# The color appearance of three-dimensional, curved, transparent objects

**DOI:** 10.1101/2019.12.30.891341

**Authors:** Robert Ennis, Katja Doerschner

## Abstract

Studies on the perceived color of transparent objects have elucidated potential mechanisms but have mainly focused on flat filters that overlay a flat background. However, studies with flat filters have not captured all aspects of physical transparency, such as caustics, specular reflections/highlights, and shadows. Here, we investigate color matching experiments with three-dimensional transparent objects for different matching stimuli: a uniform patch and a flat filter overlaying a variegated background. Two different instructions were given to observers: change the color of the matching stimulus until it has the same color as the transparent object (for the patch and flat filter) or until it has the same color as the dye that was used to tint the transparent object (for the patch). Regardless of instruction or matching element, observers match the mean chromaticity of the glass object, but the luminance of matches depends on the backgrounds of the test image and the matching element, indicating that a color constancy-esque discounting operation is at work. We applied three models from flat filter studies to see if they generalize to our stimuli: the convergence model and the ratio of either the means (RMC) or standard deviations (RSD) of cone excitations. The convergence model does not generalize to our stimuli, but the RMC generalizes to a wider range of stimuli than the RSD. However, there is an edge case where RMC also breaks down and there may be additional features that trade-off with RMC when observers match the color of thick, curved transparent objects.

## 1 Introduction

Transparent objects allow some light to pass through their bodies, as opposed to opaque objects, which only absorb and reflect light. The conditions that lead to the perception of flat transparent filters and the factors that determine the perceived color of a flat filter have been extensively studied. Helmholtz and Koffka wrote about the perception of transparency in their books, where Helmholtz described the percept of transparency as “…a transparent colored veil … spread over the field …” and Koffka referred to it as “color scission”, in which the visual system can assign more than one color to the same image region, effectively splitting that region into layers and perceiving a colored filter overlaying another colored object. However, it was Metelli’s work [55, 56] that has had a more lasting imapct on the study of perceived transparency, resulting in the classical Metelli or episcotister model of transparency, which is still used in an altered form in computer graphics, where it is known as alpha blending [29]. Metelli used an episcotister, a device that rotates a sliced disk in front of a complete and stationary disk. In some of his experiments, the stationary disk was painted such that the left half had one gray value and the right half had another, creating a vertical split in luminance down the middle (a bipartite field) that the sliced rotating disk overlaid. The sliced rotating disk was painted with one uniform gray value. When the sliced disk is spun fast enough, optical mixing of the reflected light leads to the appearance of a solid, non-rotating transparent filter in front of the bipartite field. Metelli determined a relationship between the reflectances of the fields of the two disks that predicts the amount of perceived transparency. Building on this, Beck et al. suggested that it is rather the relationships amongst the lightness percepts, instead of luminance or reflectance, that determines perceived transparency, which Metelli later adopted [57]. In fact, for the episcotister stimulus, whether a surface will appear transparent or not can be predicted by the relative luminance relationships between adjacent patches, without regard to their absolute luminance [7, 9]. In the episcotister model, these relationships must occur at the well-known X-junctions, where they determine perceived depth order [56, 11, 13, 7, 33, 42, 59, 67, 9, 53, 77, 49, 50, 15]. However, the episcotister model is not predictive of changes in transparency perception as other factors of the stimulus are varied, such as the mean luminance. For example, Singh and Anderson showed that perceived transparency varies with changes in mean luminance, which Metelli’s model says should not happen, and they concluded that it was rather Michelson contrast that determined perceived transparency. Although, it has been suggested that Singh and Anderson’s model does not incorporate the psychophysical “principle of independence of effects” for flat filters [53]. Regardless, there are situations which give rise to the perception of transparency when the established luminance relationships at X-junctions are broken. Plus, transparency is even perceived without X-junctions [82, 74], so the episcotister model of transparency perception has seen less application in modern perceptual transparency research, although the episcotister itself continues to be used as a stimulus in studies.

All of the episcotister studies mentioned above were done in the achromatic domain. If the scene and filters are chromatic, then D’Zmura et al. found that if one considers the pixels in some extended and continuous region of a Mondrian and one then shifts the chromaticity of all of those pixels in the same direction and by the same amount, then one will perceive a flat colored transparent filter overlying the Mondrian. A similar effect is obtained when forcing the chromaticity of those pixels to converge on the same color and a mixture of both techniques also works. This effect is especially curious, since transparent objects will always reduce the luminance in the region of the image that they filter, relative to when the transparency is absent, since unless they are perfectly transparent, they will always absorb and/or reflect some of the incident light. However, the technique of Dzmura, et al. does not change luminance, so the perceived transparency that their technique generates is physically impossible [19, 14, 20]. Regardless, their model is predictive of the colors that observers set when they can adjust the color in a restricted portion of a filter [14] and it is also predictive of the colors that observers set when they can adjust the color of surfaces seen through the filter [20].

However, the convergence model appears to only be applicable to the classical flat filter models that have been the primary focus of transparency perception research. For example, consider Fig 15. In this figure, we show two ways of interpreting and applying the convergence model: 1) as the most general Affine model considered in the original papers and 2) in terms of the divergence operation from vector calculus [75]. In panel A of Fig. 15, we display an example of the stimuli used in this paper: a curved glass object (a Glaven [65, 66]), sitting in a room with multicolored walls and illuminated by a spherical light source to the right and above (out of the direct view of the camera). Technical details about the analyses are contained in the “Supplemental Material”, but using these images, we applied the convergence model to judge if it can predict the color changes induced by the transparent object in our images.

In panels B and D of Fig. 15, we provide a test of the convergence model for the example image. In panel B, we show the actual filtered colors, coming from the original image in panel A as blue points in the isoluminant plane of the MB-DKL space. The red points show the result of applying the best fitting Affine transformation to the colors of the wall that are eventually filtered by the glass Glaven. Ideally, the red points and blue points should line up on top of each other and RMSE should be low, but it is at a value of 0.21 in this case, which is well above the amount of error one would like in the MB-DKL space, relative to the scaling of its axes (typically in a range of −1 to 1). Note that although in this case we are showing the data projected into the isoluminant plane, the Affine map considered for panel B was found for the full three-dimensional data, including luminance, which does not improve the fit.

In Table 1, we show the average RMSE for the best-fitting Affine maps. Note that in this table we average over fits for the full three-dimensional distributions, including luminance, and for two-dimensional isoluminant projections, since we found little difference in RMSE for both types of fit, as can be seen in their small overall standard deviation. Otherwise, the average RMSE were grouped according to color space and mask type. We tried three masks: one that captured all pixels filtered by the transparent object; one that excludes the specular highlight from the calculations; and one that excludes all specular reflections from the calculations. The specular reflections were found by putting the specular reflection component of the transparency BRDF in Mitsuba (the physically-based renderer we used (see “Methods”)) to its maximum value and putting the transparency component to zero (i.e., full absorption of all incoming light, giving a black surface). Note that we also performed the analysis described above in the CIELAB color space. The resulting RMSE values are also shown in Table 1.

**Table 1:**
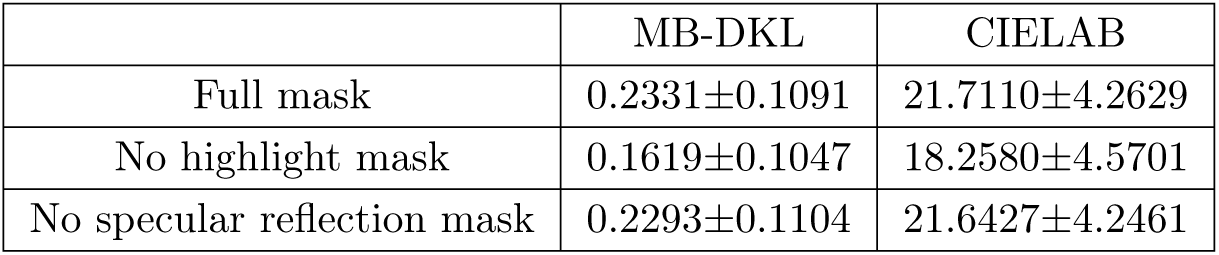
Average RMSE (±SD) for fits of the general Affine map from the original convergence model papers to our stimuli. See the main text and “Supplemental Material” for more details.

|  | MB-DKL | CIELAB |
| --- | --- | --- |
| Full mask | 0.2331±0.1091 | 21.7110±4.2629 |
| No highlight mask | 0.1619±0.1047 | 18.2580±4.5701 |
| No specular reflection mask | 0.2293±0.1104 | 21.6427±4.2461 |

Panel D of Fig. 15 shows the results of the divergence analysis. Divergence specifies at each point the amount that vectors are all moving in the same direction and radiating out from that point (i.e., diverge) or the amount that they are all pointing inwards towards that point (i.e., converge). The case in between is that all vectors are parallel to each other. Since divergence is not a single quantity, but a function that maps a scalar to each point in the space of interest, we show it as a gray-scale heatmap in panel G, where white points (> 0) denote divergence, black points (< 0) denote convergence, and gray points denote that the vectors are oriented parallel to each other (= 0). It can be seen that there is no clear region with substantial black, which would indicate a point of convergence that would be parsimonious with the previously described convergence model. Rather, we find a salt-and-pepper scattering of values with no clear structure and this is the same result for all the images we tested. Please note that this divergence analysis was only performed on the chromaticity components of the vector fields (i.e., after they had been projected into the isoluminant plane).

Considering the above, it seems that the convergence model is not generalizable to curved transparent objects that exhibit specular reflections, shadows, and caustics. However, there are other models of perceptual transparency that could generalize from the flat filter case to the full three-dimensional case. There are more recent attempts to investigate perceptual transparency with a physically-based model [8, 58]. The model includes internal refraction and filtering effects on luminance. These studies have followed two closely related paths that have come to somewhat different conclusions. The investigations of Khang, Robilotto, and Zaidi [43, 44, 73, 72] have found that observers use a measure of contrast to detect and match a flat filter [72, 73, 43], since the mean color in the filtered region is insufficient to predict observer matches. In fact, when asked to match the color of two filters, each placed under a different illuminant, observers match the ratio of mean cone excitations between the filtered and unfiltered region (RMC) [43]. Khang and Zaidi also found that observers do not match the chromaticity difference between filtered and unfiltered regions, but they do match the ratio of mean chromatic contrast. Here, we focus on the RMC that directly involves cone activations, since this is more directly comparable with other research described below. The ratios of mean cone excitations are similar to, but not exactly the same as, mean spatial cone excitation ratios (MCER), which have been extensively studied with respect to color constancy [30, 61, 60, 62, 31, 32]. Actually, the work of Ripamonti and Westland [83, 71] found MCER capable of predicting the strength of perceived transparency. Essentially, Khang, Robilotto, and Zaidi’s investigations have found that observers are capable of making veridical identifications of filters across different illuminants and backgrounds [44], achieving color constancy, and that observers match a quantity that is related to spatial variation and the change in cone signals between the filtered and unfiltered regions. From this perspective, flat filters act similarly to spotlights [45, 18, 47]. The investigations of Faul and Ekroll, Faul and Ekroll, Faul and Ekroll, Faul and Falkenberg made a detailed investigation of the same physical model [8, 58] and devised a way to transfrom the physical parameters of the model into various perceptual quantities, such as the hue and saturation of the filter, provided that certain simplifying assumptions are adopted [26]. Their model suggested that observers would match the ratios of the standard deviations of cone excitations between the filtered and unfiltered regions (RSD), unlike the RMC mentioned above, and their psychophysical experiments supported this conclusion. Using their perceptual model of filter color, they also found that when matching filters, each placed under a different illuminant, observers make a match that lies approximately halfway between a proximal match and a perfect color constancy match [28].

However, while the results of studies with flat filters have provided a great deal of information about how transparency is perceived, they do not incorporate other factors that arise when considering three-dimensional, curved, transparent objects. The physical process of transparency in a variegated three-dimensional environment gives rise to shadows, caustics, specular highlights, subtle reflections of the surrounding environment, and a tinted and refracted (i.e., distorted) view of surfaces that lie beyond the transparent object’s body. Shadows, caustics, specular highlights, and subtle reflections of the environment are not present in studies with flat filters and all could play a role in the color appearance of a transparent object. For example, caustics are unique to more translucent and transparent objects and are rather complex, being determined by the geometry of the illumination, the shape and refractive index of the transmitting object, and the shape of the surface which will receive these focused regions of light. Caustics may be too complex to serve as a cue to the structure of the illumination, but their color is heavily influenced by the transmission and absorption spectrum of the transmitting body, so they may play a role in the color appearance of the transmitting body.

In this paper, we investigate what information observers use when judging the color of 3-D, curved, transparent objects. We present here variations of color matching experiments that attempt to answer the question: “What is it that makes a red glass look red?” and we specifically test whether the RMC or RSD is capable of predicting observer matches.

## 2 Materials and methods

### 2.1 Scene rendering

To investigate the features that observers might be using when judging the color of 3-D transparent objects, we used the multispectral and physically-based Mitsuba renderer (version 0.6.0, from Github; git commit: 06340ccbb3f4) to generate 400px × 400px images of a square room with a glass object, called a Glaven [65, 66], and a small spherical light. The spherical light (not a point light source!) was placed above and to the right of the Glaven, outside the field of view of the camera, and it created a noticeable highlight on the top right of the Glaven. One example stimulus is in Fig. 2 and more examples can be found in Fig. 3. There are two things to notice: first, the Glaven is hollow inside, but the walls have a thickness to them, and second, if one looks closely, they can see two highlights overlapping each other. This is due to internal reflection between the inner and outer side of the wall of the Glaven. The highlight is tinged with the color of the transparent object, due to total internal reflection at the inner wall boundary, which is not something seen with glossy objects.

**Figure 1:**
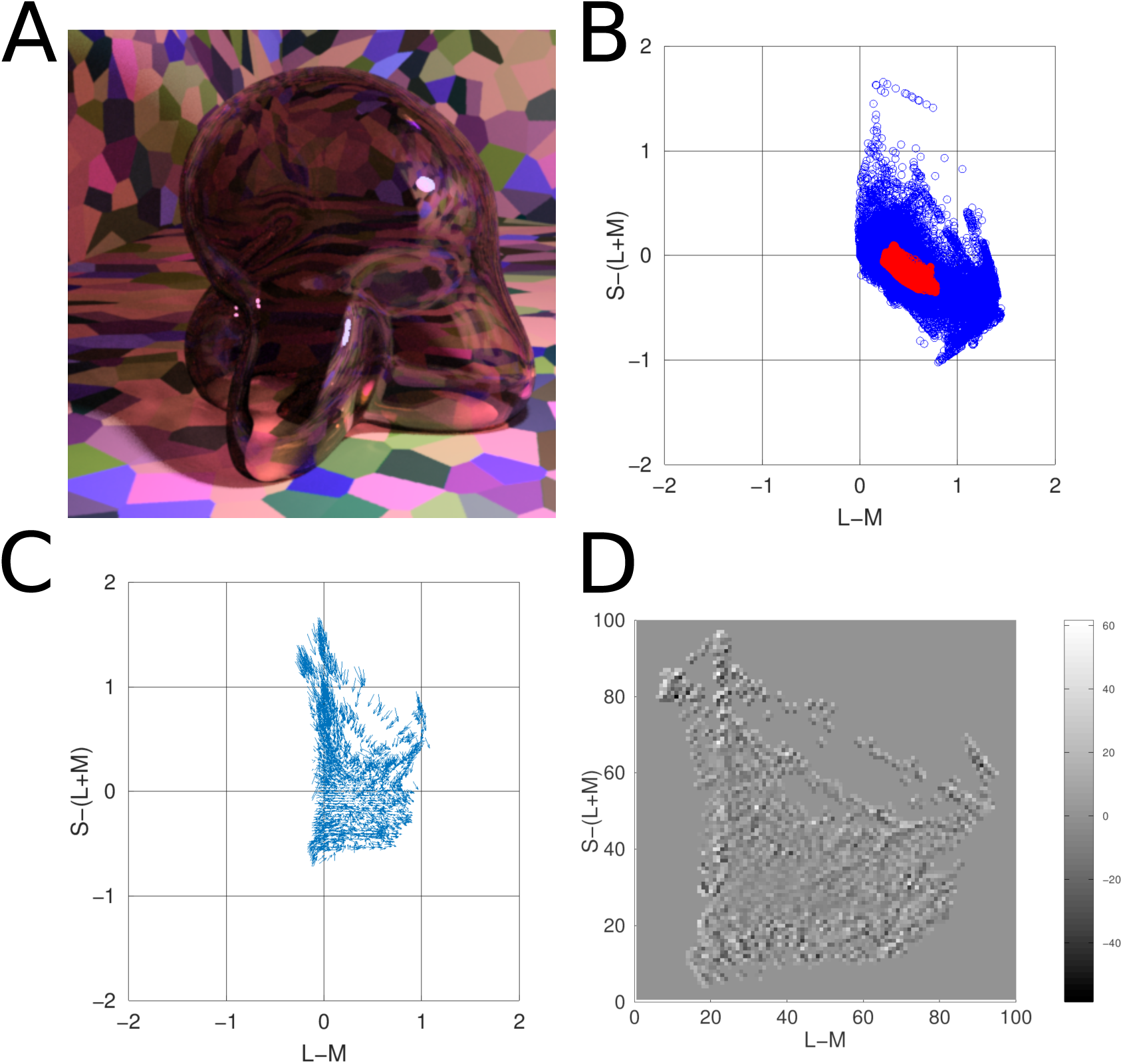
Results of applying two forms of the convergence model to the stimuli used in this paper. Panel A) One of the actual stimuli used in the paper: a colored, curved, glass object (a Glaven), placed in a room with multicolored walls and illuminated by a spherical light source to the right and above (out of view of the camera). Panel B) The isoluminant plane of the MB-DKL color space, where the colors of the pixels that contain the colors directly filtered by the glass Glaven are plotted in blue. The red pixels show the result of applying the best-fitting Affine transformation to the colors of the unfiltered pixels in panel B of Supplemental Fig. 1. Panel C) A vector field in the isoluminant plane of the MB-DKL color space. Each arrow connects the color from a given pixel in the unfiltered image of panel B of Supplemental Fig. 1 (vector base) to the color of that same pixel after being filtered by the transparent object in panel A (vector tip). Please note that this vector field is an interpolated one, which was necessary for the divergence analysis described later, and the divergence analysis was only computed for the isoluminant-projected distribution shown. Please see the “Supplemental Material” for more details. Also, note that the scaling of the vectors is arbitrary, since we plotted the data with MATLAB’s quiver function (R2018b; Mathworks, Inc.; Natick, Mass., USA). Panel D) A heatmap showing the divergence values computed for the interpolated, isoluminant vector field shown in panel C. White (> 0) indicates divergence, black (< 0) indicates convergence, and gray indicates parallel vectors in the local neighborhood of that location (= 0). A salt-and-pepper pattern with no clear region of convergence or divergence can be seen.

**Figure 2:**
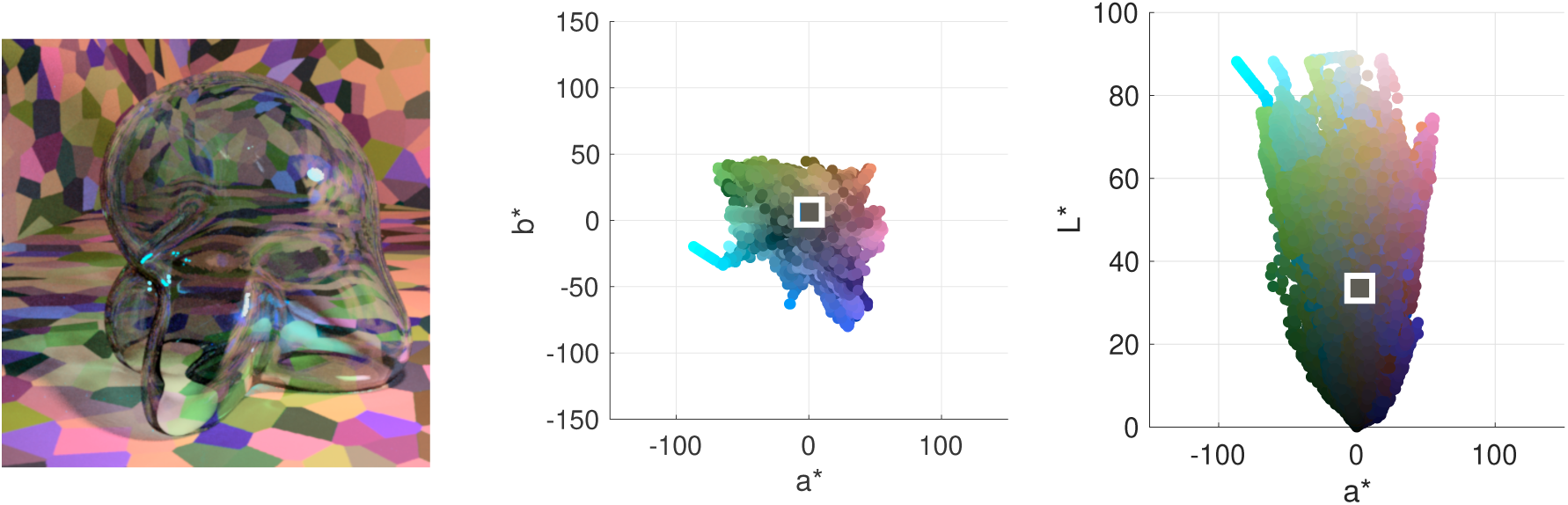
(Left) A close up example of one of our stimuli: a glass Glaven with the blue Munsell high transmission distribution under the white illumination. (Middle) The CIELAB coordinates in the (a*, b*) plane for the pixels from the filtered region of the image (i.e., the region covered by the Glaven). The points are colored with their corresponding RGB coordinates. The white box is centered on the average of the distribution. (Right) The CIELAB coordinates in the (a*, L*) plane for the pixels from the same filtered region of the image. The points are again colored with their corresponding RGB coordinates and the white box is centered on the average of the distribution.

**Figure 3:**
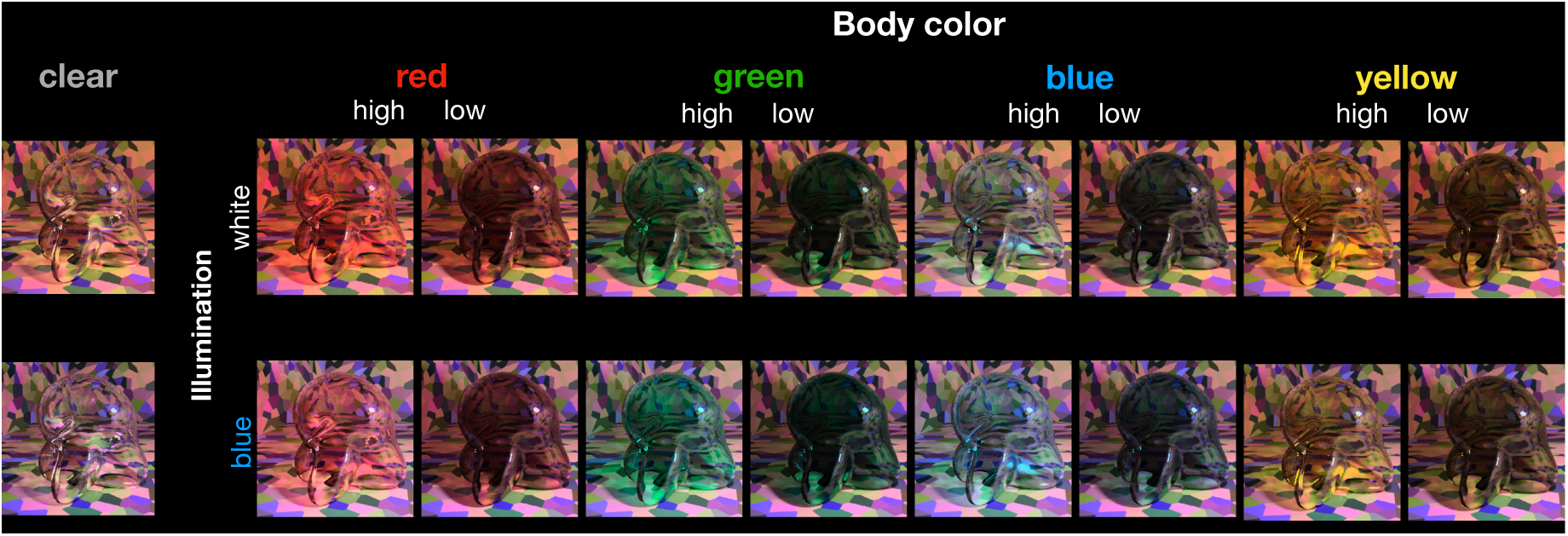
Our initial set of test images. The 16 images containing the red, green, blue, and yellow transparent Glavens were presented to observers during the main color matching experiments. Two transmission levels were tested (high and low). The high transmission condition was simply the original Munsell reflectance distribution that we gave to Mitsuba and the low transmission condition was made by scaling the distribution down by a constant factor of 0.68. The Glavens were rendered under two different illuminants: a blue and a white illuminant, each from the daylight locus. The two images at the left, showing the clear Glavens, illustrate that the illuminant can have a subtle effect on the color appearance of curved, transparent objects that are highly transmissive. See the text in the Methods for specific details about the rendering procedure.

The 3-D configuration of the scene was created in Blender (v2.79b) and was exported to Collada DAE format. The DAE file was then translated into a serialized binary file that Mitsuba can efficiently process, using Mitsuba’s mtsutil program. The mtsutil program produces an XML file that specifies to Mitsuba the scene layout, the properties of objects in the scene, and the details of the simulated camera system that “takes the image” during the rendering process. The XML file was edited so that the Glaven had Mitsuba’s smooth dielectric BXDF with the default properties that make it act like glass. We then used a small script, written in R (v3.6.1), to produce variations of this XML file that used different spectral distributions for the illumination, different transmission distributions for the dielectric BXDF, and to give the wall either a uniform Lambertian BXDF with a flat reflectance distribution (i.e., white walls) or to texture the wall with a multicolored RGB Voronoi pattern with a Lambertian BXDF. We also found that we could only achieve acceptable rendering of caustics with the small, spherical light source and with the Metropolis Light Transport integrator in combination with the Independent sampler running at 1000 samples per pixel.

The color of the glass Glaven was varied by providing different Munsell spectral reflectance distributions to the spectral transmission and spectral reflectance parameters of the dielectric BXDF. Four Munsell spectral reflectance distributions were chosen from the database provided by the Computational Spectral Imaging group at the University of Eastern Finland [39]. They corresponded to the chips listed in Table 2.

**Table 2:** The chips whose spectral reflectance distributions were used as the spectral transmittance distributions of the glass Glaven when it was rendered with Mitsuba.

|  |  |
| --- | --- |
| Red | 2.5R 7/2 |
| Green | 5G 2.5/2 |
| Blue | 5B 4/1 |
| Yellow | 10Y 5/1 |

The Munsell reflectances were chosen such that if they were used as the spectral reflectance distributions for a piece of paper and these hypothetical papers were measured under an equal-energy white, then their chromaticity coordinates would lie close to the cardinal axes of the MacLeod-Boynton-Derrington-Krauskopf-Lennie color space (MB-DKL) [52, 17] (Standard procedures for converting between linear RGB coordinates and the MB-DKL space are documented elsewhere [86, 37]). These axes selectively stimulate the S-(L+M) and the L-M mechanisms [16]. We also made four more spectral distributions that were scaled versions (scaling factor = 0.68) of the Munsell spectral reflectance distributions that were just described. These scaled versions, when used as the spectral transmission and spectral reflectance distributions of the glass BXDF, result in a darker and more saturated color appearance for the glass. Color coordinates for these Munsell reflectances and the illuminants (described below) in the CIE1931 xyY space are depicted in Fig. 4.

**Figure 4:**
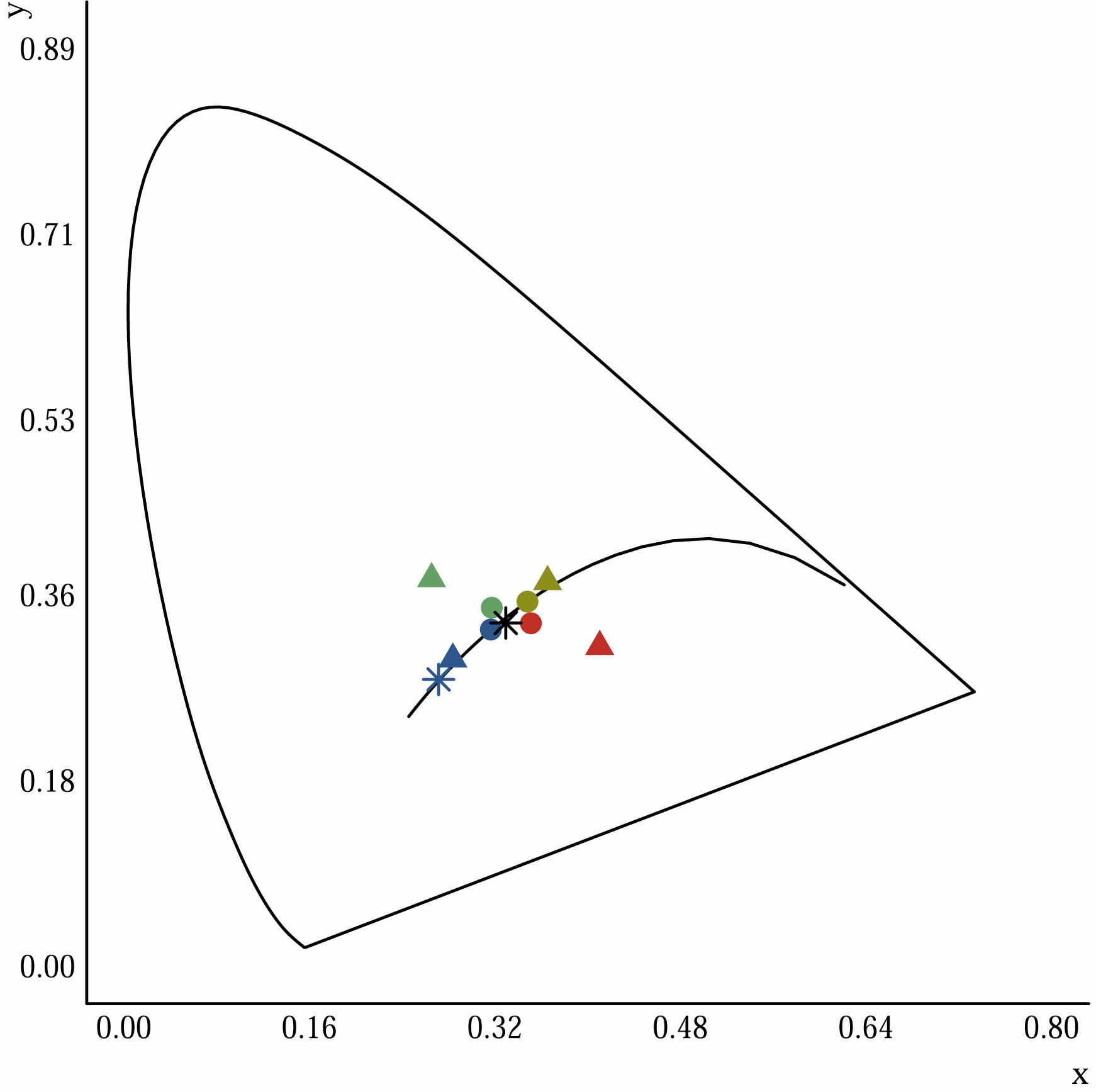
Representation of illuminants, stimulus transmission distributions, and the matching filter distributions in the CIE1931 xyY chromaticity diagram. The colored stars denote the blue (blue symbol) and white (black symbol) illuminants that were given to Mitsuba when rendering our scenes. The colored disks denote the chromaticity coordinates of the four Munsell transmission distributions (red, green, blue, and yellow) that were used when rendering the Glavens. They correspond to a flat filter, lying above a white surface and imaged under a neutral illuminant. Since changing the intensity of a distribution does not change its position in the chromaticity diagram, the high and low transmission Munsell distributions plot directly on top of each other and so only four of the eight disks can be seen. The triangles denote the four Munsell distributions (red, green, blue, and yellow) that were used as endpoints of two filter transmission axes (red-green and blue-yellow). During experiments described in the “Experimental paradigm and stimulus presentation” section below, observers could vary the transmission distribution of a matching filter through linear combinations of the two filter transmission axes.

For the multi-colored Voronoi background, all eight of these spectral distributions were rendered under a blue and a white illuminant, giving 16 combinations in total. (Note that the images with the white walls were rendered under a neutral illuminant metameric to D65.) The spectral distributions of these illuminants were obtained from prior work that used hyperspectral measurements of real scenes [23]. Briefly, the illuminants were measured under a JUST Normlicht LED box by placing a PhotoResearch white reference in the box at approximately a 45° angle and placing the line of sight of the hyperspectral camera roughly perpendicular to the white reference surface. The two chosen illuminants lie along the daylight locus and were created from linear combinations of the blue and yellow illuminants taken from the just referenced hyperspectral measurements. The spectral distributions of the illuminants were first scaled down by 50%, because otherwise highlights would be burnt-out and map to maximum white (i.e., all RGB values in the highlight at 100%). These dimmer illuminants were then used in the next step: each illuminant was slightly desaturated by scaling it down to 70% of its maximum and mixing it with 30% of the other distribution. For example, a relatively slightly desaturated blue illuminant was produced by scaling it down by 70% and adding the spectral distribution of the yellow illuminant after it was scaled down to 30% of its maximum. This was done because while highly saturated illuminants produce physically-reasonable images, they will have a reduced variance of hues and the scenes look rather artifical. The two resulting illuminants that were used for rendering were blue and white in appearance and were on the daylight locus. This was acceptable, since we were not performing a rigorous color constancy study and were mainly interested in having some extra variety in the stimuli.

Our version of Mitsuba was configured to render with multispectral data, rendering visible light in the range of 360nm to 830nm in 47 equally spaced wavelength bands. The distributions that we provided as input to Mitsuba were specified in the range of 380nm to 830nm and were sampled either in 400 equally spaced wavelength bands (Munsell reflectances) or in wavelength bands that were on average ∼ 1.12nm wide (illuminant distributions), which is due to the construction of our hyperspectral camera. When provided with this data, Mitsuba interpolates it and resamples it at 360nm to 830nm in 47 equally spaced wavelength bands, filling in zeros where no data was provided by the user. This was more than sufficient for our purposes and produced realistic and pleasing images.

The Voronoi background seen in Fig. 3 was produced with a modified version of an OpenGL fragment shader created by Íñigo Quílez and found on ShaderToy [68]. The Voronoi algorithm determined not only the shape of each element in the texture, but also its color, which was sampled from multiplications of 3 base colors, where the intensity of each base color was modulated according to a cosine that was sampled with the ID of each element, as determined by the Voronoi algorithm, and the specific trio of base colors was also determined by the ID as well. We wrote a small OpenGL-based program in Rust (v1.30) [54, 4] that used the shader to produce a large 1024px × 1024px RGB Voronoi texture that could be fed into Mitsuba to act as a wall texture.

We also rendered the scene with a Lambertian background that had a flat reflectance distribution with 100% reflectance, giving the appearance of white walls. The uniformly reflecting Lambertian BXDF is a feature provided by Mitsuba, so we only had to specify the parameters in the XML scene file. We did not provide any external spectral data for the white walls. Note that the images with the white walls were rendered under a neutral illuminant that was metameric to D65. The reason for this is given at the beginning of the “Experimental paradigm and stimulus presentation” section below.

The final render output of Mitsuba, in our case, was an HDR OpenEXR image containing linear RGB data in 16-bit floating point format. Mitsuba uses the standard CIE1931 color matching functions [84] and the standard sRGB specifications [3] (in particular, the ITU-R Rec. BT. 709-3 primaries [2] with a D65 white point) to convert from the multispectral render output to linear RGB coordinates. Mitsuba’s built-in Reinhard tonemapper [69], provided by the mtsutil program, was used to convert these outputs to 8-bit RGB images in a PNG format. The following parameters were provided to the program: multiplier = 0.8, gamma = 1.8, key = 0.8, and burn = 0.1. All other parameters were left at their default values and we found this combination to be best at preventing burn-out of highlights.

In total, for the main experiments described here, we had 24 images: 8 with the Voronoi background and the four original Munsell distributions (4 for each illuminant), 8 with the Voronoi background and the four scaled Munsell distributions, 4 with the white background and the four original Munsell distributions, and 4 with the white background and the four scaled Munsell distributions.

### 2.2 Experimental paradigm and stimulus presentation

The task of observers was to change the color of a matching element until it appeared to have the same color as the glass Glaven shown in the test image. The Voronoi background images and the white wall background images were tested in separate experiments. The white wall images were tested to see if observers perform some kind of color constancy-esque discounting operation when they make their matches. Essentially, is the color of a transparent object determined by what the object would look like if it were placed against a white background under a neutral illuminant?

Prior to each experiment, observers adapted to the mid-gray of our monitor for one minute. A beep then indicated the start of the experiment and one of the images corresponding to the specific test set (Voronoi or white wall) was presented at the center of the monitor. The images and the matching elements were always presented against a background that was the mid-gray of the monitor and the images were always presented in a randomized order. Five matches were made for each image, resulting in 80 trials for the Voronoi set of stimuli and 40 for the white wall set of stimuli. We used two different matching elements in separate experiments: a 50 pixel × 50 pixel uniform patch or a 256 pixel × 256 pixel image of a flat transparent filter lying above an achromatic Voronoi background. The flat transparent filter was rendered according to an equation provided by Khang and Zaidi that is similar in result to the formulation given by Faul and Ekroll, Faul and Ekroll, Faul and Ekroll, Faul and Falkenberg. See Fig. 5 for examples of the matching elements. The Voronoi background was generated using the same Rust program that was used to make the Voronoi background in the rendered scenes, but it was altered to only produce various shades of gray. The resulting image was 256px × 256px.

**Figure 5:**
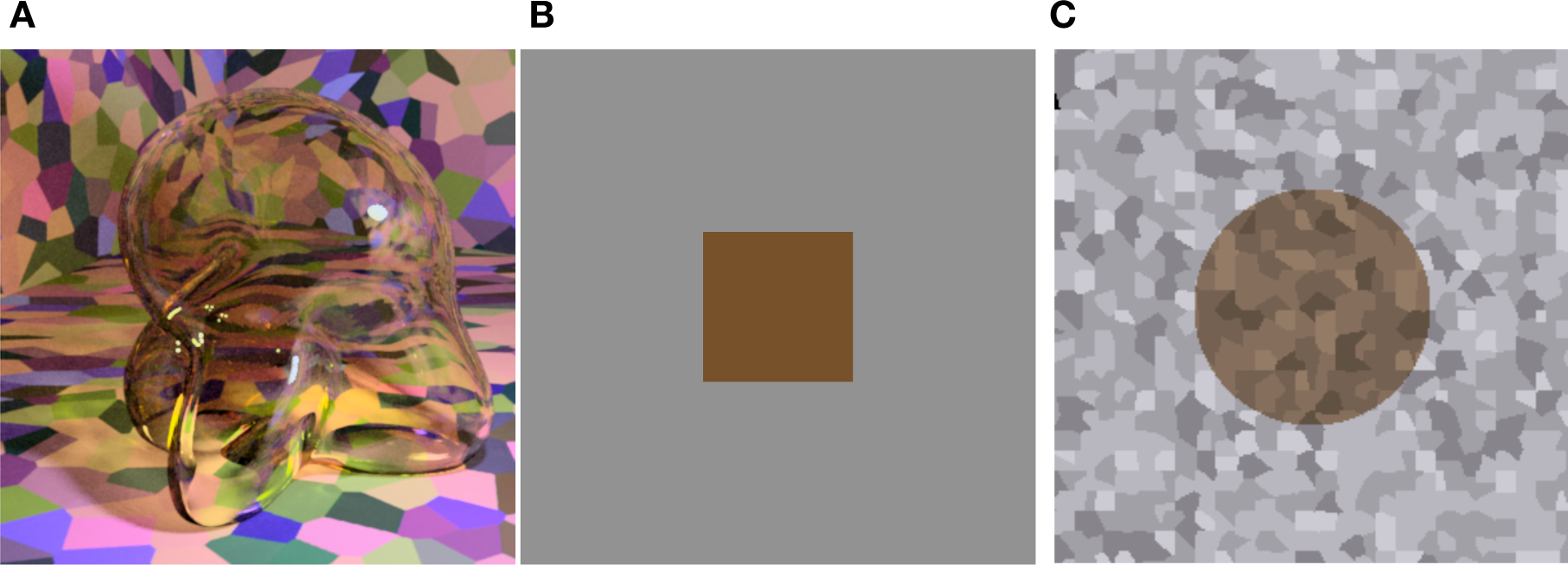
Examples of the stimuli that observers used when making a color match to the transparent Glaven. (A) An example stimulus to which an observer would make a match. (B) Observers could adjust the color of a simple patch. Shown is the average setting of observers in “proximal match” experiment for the glass Glaven shown in panel A. (C) Observers could adjust the color of a flat transparant filter, lying above an achromatic Voronoi background. Shown is the average setting of observers in our experiment for the glass Glaven shown in panel A. See the text in the “Methods” section for specific details about how the matching stimuli were created and how observers controlled the color of the matching stimuli.

We tested two different sets of instructions for the uniform patch matching element in separate experiments and one set of instructions for the flat transparent filter in an additional experiment. In the first experiment with the uniform patch, observers were asked to adjust the color of the patch until it appeared to have the same color as the glass Glaven (“proximal match”). This was similar to a proximal stimulus match in color constancy research. In a follow-up experiment, observers were asked to adjust the color of the patch until it appeared to have the same dye that was used to tint the glass Glaven (“dye match”). This was similar to asking observers to make a paper match in color constancy experiments. In the flat transparent filter experiments, observers were asked to “change the color of the glass filter until it appeared to have the same color as the glass Glaven”.

In all experiments, observers adjusted the color of the matching element by moving the mouse. The mouse cursor was hidden from view during the experiment. As an example of how mouse position was mapped to color, we can consider the uniform patch stimulus, where its color was specified in the MB-DKL space. When the position of the cursor was at the center of the screen, the uniform patch was mid-gray. When the cursor was moved left, the color of the patch was shifted a proportional amount along the green direction of the L-M cardinal axis of the MB-DKL space. When the cursor was moved right, the color of the patch was shifted a proportional amount along the red direction of the L-M cardinal axis. Up and down were similarly mapped to the blue and yellow directions of the S-(L+M) cardinal axis. With this setup, moving the mouse around in a circle at a given radius from the center of the screen would modulate the uniform patch through all colors on a hue circle at a proportional radius in MB-DKL space. The luminance of the patch could be adjusted by pressing the left mouse button to make it darker and the right mouse button to make it brighter.

The normalized mouse position was computed by subtracting the mouse coordinates for the center of the monitor from the current mouse position and dividing the x-component by half the width of the monitor (specified in pixels) and the y-component by half the height of the monitor (specified in pixels).

In the case of the flat transparent filter, the achromatic Voronoi image was loaded into MATLAB and a circular region at the center of the image with a radius of 60 pixels was processed according to the filter formula in Khang and Zaidi. Mouse position was now mapped to linear combinations of four Munsell spectral reflectance distributions that defined the transmission distribution component of the filter equation. These four Munsell distributions were also taken from the Computational Spectral Imaging group at the University of Eastern Finland [39] and they corresponded to the chips listed in Table 3.

**Table 3:** The Munsell chips whose spectral reflectance distributions were used in the observer-controlled linear combination that determined the spectral transmittance distribution of the flat filter matching element.

|  |  |
| --- | --- |
| Red | 7.5RP 6/8 |
| Green | 10G 7/8 |
| Blue | 7.5B 2.5/2 |
| Yellow | 7.5Y 5/2 |

These four Munsell distributions were all normalized, such that their maxima were equal to 1, and they were taken in pairs as the endpoints of two axes, one pair defining a “red-green” variation and the other defining an orthogonal “blue-yellow” variation. Now, moving the mouse would vary the shape of the transmission distribution of the filter and clicking the mouse buttons would vary the overall transmittance, which would change the apparent opacity or “lightness” of the filter. The changes were enacted according to the following equation:

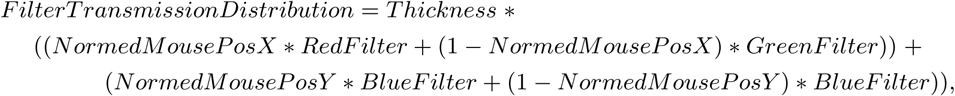

where *Thickness* was increased or decreased according to left and right mouse clicks, respectively. This scheme produces a transmission distribution color space that is analogous to the MB-DKL color space and moving the mouse in a circle at a given radius from the center of the monitor would modulate the color of the filter through all the hues.

In all experiments, when observers were satisfied with their match, pressing the middle mouse button would clear the screen, save their data, and prepare the next trial. Before each trial, observers re-adapted to the mid-gray of the monitor for two seconds.

### 2.3 Monitors

Stimuli for the Voronoi background stimuli (“dye match”, “proximal match”, and “filter match”) were displayed on a 10-bit EIZO ColorEdge CG23W monitor (EIZO Corporation; Hakusan, Japan) using Psychtoolbox (SVN revision 8643) [12, 64, 46] via the MATLAB environment, R2014b (Mathworks, Inc.; Natick, Mass., USA). The computer that was connected to the EIZO was a Dell Precision T1700, running Microsoft Windows 7 Professional edition SP1 (64-bit) (Microsoft Corporation; Redmond, Wash., USA) with an Nvidia Quadro K620 graphics card (Nvidia Corporation; Santa Clara, Calif., USA) controlled by version 347.52 of the Nvidia drivers.

For the experiment with the white walls (filter match), stimuli were displayed on a SONY PVM2541-A OLED (Sony Corporation; Tokyo, Japan) via Psychtoolbox (SVN revision 9641) [12, 64, 46] using the MATLAB environment, R2018b (Mathworks, Inc.; Natick, Mass., USA). Although the SONY OLED is a 10-bit monitor, it requires special hardware to be used in this mode, so our SONY OLED was running in 8-bit mode, but this was found to not have an effect on the final results of our experiments. The computer that was connected to the SONY OLED was an HP Pavilion 595 (HP, Inc.; Palo Alto, Calif., USA), running Microsoft Windows 10 Home edition (64-bit) with an Nvidia Geforce GTX 1050 Ti controlled by version 398.36 of the Nvidia drivers.

Both computers were disconnected from the internet to prevent automatic updates from potentially changing their behaviour. Both monitors were calibrated using a Konica-Minolta CS2000-A via standard procedures documented elsewhere [86, 37]. In particular, the calibrations were used to ensure that our stimuli could be accurately reproduced by the gamuts of both monitors and to calculate CIELAB [1, 84] representations, and LMS cone excitations [79, 78] for our stimuli (explained in further detail below, in the “Color space conversion” subsection). The properties of the Eizo monitor in the CIE1931 xyY colorspace were as follows: red phosphor (x: 0.6733, y: 0.3088, Y: 37.223), green phosphor (x: 0.2103, y: 0.6869, Y: 86.659), and blue phosphor (x: 0.1564, y: 0.0554, Y: 7.9206). The properties of the SONY OLED monitor in the CIE1931 xyY colorspace were as follows: red phosphor (x: 0.6694, y: 0.3235, Y: 37.675), green phosphor (x: 0.1924, y: 0.719, Y: 82.7), and blue phosphor (x: 0.1436, y: 0.0539, Y: 9.6269).

### 2.4 Observers

We had different groups of observers in each of the experiments: 6 that adjusted the uniform patch with the “proximal match” instructions and the multi-colored Voronoi background, 9 that adjusted the uniform patch with the “dye match” instructions and the multi-colored Voronoi background, 5 that adjusted the flat transparent filter and the multi-colored Voronoi background, and 10 that adjusted the flat transparent filter and the white walls. Three observers also ran in a control experiment discussed in more detail later in this manuscript. One of the observers in this control experiment was one of the authors, RE.

All observers were in the age range of 20-30 years old, so yellowing of the macular pigment cannot be considered a significant contribution to our results, and they were all recruited from the student population at the Justus-Liebig University in Giessen. The observers were naïve to the purpose of the experiment. All observers had normal or corrected to normal visual acuity. All observers were checked for color deficiency using the Isihara color plates [40] and no observers were excluded based on its criteria. Observers were paid for their participation in the experiments. All observers gave written informed consent in accordance with the Code of Ethics of the World Medical Association (Declaration of Helsinki) for experiments involving humans. The experiments were approved by the local ethics committee LEK 2015-0021.

### 2.5 Analysis

#### 2.5.1 Color space conversion

For analysis, we converted our rendered images and the settings that observers made with the matching elements to the CIELAB color space [1, 84] and LMS cone excitations [79, 78]. The conversions were done with routines written in the Go programming language (v1.13.3) [6]. Essentially, the routines convert the RGB values at each pixel to the corresponding CIELAB or LMS coordinates, using the equations described below, and returns a CIELAB “image” or LMS “image” respectively.

The CIELAB color space was developed to be a perceptually uniform color space, meaning that movement in any direction at any point in the space by a unit amount should correspond to a step size of 1 JND. One feature of the space that assists with this goal is accounting for adaptation to the illuminant, whose CIE1931 XYZ coordinates play a role in the transformation equations that define the CIELAB space. While the original CIELAB definition does not achieve the goal of complete perceptual uniformity, it is close enough for our our purposes here. The CIELAB space has three color axes which correspond to red-green, blue-yellow, and light-dark variations. The equations that map a color to the CIELAB space require the CIE1931 XYZ coordinates of the color and the CIE1931 XYZ coordinates of the illuminant (also known as the “reference white point”). The equations are as follows:

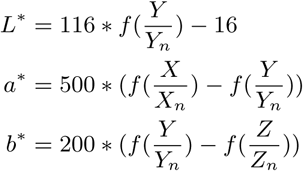

and

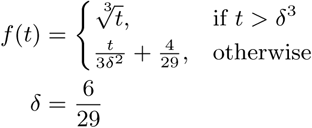

where *X*_*n*_, *Y*_*n*_, and *Z*_*n*_ are the CIE1931 XYZ coordinates of the reference white point and *X, Y*, and *Z* are the CIE1931 XYZ coordinates of the color of interest. L* aims to be a perceptually uniform scale for luminance, a* aims to be a perceptually uniform scale for red-green variations, and b* aims to be a perceptually uniform scale for blue-yellow variations.

When converting our images to the CIELAB color space, the chromaticity coordinates of the maximum white of the monitor were used as the reference white point.

The LMS excitations were computed using the calibration data of the monitors. Briefly, an {*L, M, S*} triplet can be computed from an {*R, G, B*} triplet in the range of [0, 1] with knowledge of the spectral distributions emitted from the three primaries when they are at their maximum intensity. Once these were obtained, we used the 2° LMS cone spectral sensitivity functions [79, 78] to calculate the {*L, M, S*} excitations for the three primaries. Then, provided that the primaries do not change in their properties as their intensity changes, one can take a given {*R, G, B*} triplet and scale and sum the maximum {*L, M, S*} excitations accordingly to get the total {*L, M, S*} excitation. For example, if the {*R, G, B*} triplet = {0.5, 0.2, 0.5}, then the total {*L, M, S*} excitation = {0.5 · (*L*_*R*_ + *L*_*G*_ + *L*_*B*_), 0.2 · (*M*_*R*_ + *M*_*G*_ + *M*_*B*_), 0.5 · (*S*_*R*_ + *S*_*G*_ + *S*_*B*_)}.

#### 2.5.2 Image statistics

We assessed how well different image statistics could predict the matches of observers. These statistics were computed on the CIELAB and LMS color distributions of the pixels filtered by the glass Glaven. The statistics were computed with a program written in the Go programming language (v1.13.3) [6] that used the color space conversion routines mentioned in the previous section.

Essentially, the Glaven was segmented from the image and the analysis was run on this segmented region. This included the highlight and some of the caustics. For the uniform patch matching element, we considered the following statistic:

- Average along the three axes of the CIELAB space for the filtered region of the image

For the flat filter matching element, we considered an expanded set of statistics:

- Average along the three axes of the CIELAB space for the filtered region of the image
- The color of the brightest pixel for the flat filter, when represented in CIELAB space (a.k.a., “White Point”)
- Ratio of mean cone excitations for filtered and unfiltered regions of the image, as suggested by Khang and Zaidi
- Ratio of the standard deviation of cone excitations for filtered and unfiltered regions of the image, as suggested by Faul and Ekroll, Faul and Ekroll

The ratio of mean cone excitations is computed by segmenting the image into filtered and unfiltered regions. In the case of the Glaven test stimuli, the filtered region is that region that is covered by the glass Glaven itself, and the unfiltered region is everything else. Similarly, for the flat filter matching stimulus, the filtered region is that covered by the simulated filter and the unfiltered region is the achromatic Voronoi background surrounding it. Then, for each of these regions, the mean excitations of the L, M, and S cones at each pixel are computed separately. Last, the ratios of these means (again, computed separately for each cone class) are calculated, with the filtered region typically in the numerator and the unfiltered region in the denominator. This results in a vector with 3 elements: [ratio L, ratio M, ratio S]. The ratio of standard deviation of cone excitations is computed in the same exact way, except the mean is replaced by the standard deviation.

The “White Point” statistic mentioned above is essentially the “brightest point on the object is most informative about albedo/surface color” rule [34, 81, 35]. Note that these statistics are computed directly on the colors in the image. When computed in this fashion, there is no preprocessing stage that performs any color constancy operations.

#### 2.5.3 Analysis of observer data

Since observer data was collected in MATLAB and saved in its MAT file format, a program written in Octave (v5.1.0) [22] was used afterward to cycle through each observer’s data and convert it to a CSV format that could be easily processed in the R programming environment (v3.6.1) [5]. For data that came from the experiments that used the flat filter matching element, the Octave program also used the saved data to re-create the flat filter that observers set for their match by passing the saved parameters for their match to a program written in the Go programming language (v1.13.3) [6]. This Go program would then compute and return the statistics mentioned above for the flat filter, using the same routines mentioned above. After these processing stages, R programs were written to analyze data and make plots. Only base R packages were used for analysis and plotting. In particular, the key statistic computed for data that came from observers was the grand mean: that is, the mean of the observers’ mean settings, and so errorbars in plots always show the standard error of the mean (SEM). In plots that show best-fit lines for data, these lines were always fit to the distribution of observer means, not to the grand mean data. In other words, they were fit to the data that were used to compute the grand mean. This was done to better capture the variance in the data via the fit. In the end, the differences in doing fits either way were negligible.

In all of our experiments, it was possible for observer settings to go out of the monitor gamut (i.e., they could make matches which had R, G, or B values that were less than 0 or greater than 1). When this happens, the monitor will either clamp the RGB values to the range of [0,1] or it presents a random color. To prevent any contamination of results, we excluded any settings that went out of gamut. This was rather rare. In the worst case, for the experiment where observers viewed the Glaven against a white wall and used the flat filter as a matching element, 93 out of the total of all 681 trials across all observers had to be excluded. Overall, the average percent of trials that had to be excluded from an experiment was 6.55% +/- 4.36% of all trials across all observers for the given experiment. This was deemed acceptable and in spite of rejecting a few trials, we were led to parsimonious conclusions across all experiments and consistent results across all observers.

#### 2.5.4 CIEDE2000 maps

It is already known from work by Giesel and Gegenfurtner and Toscani et al., Toscani et al. that when making color matches to Lambertian objects, observers use the most luminant region of the object to guide their match. To gain some perspective on which regions of the object observers might use when making a match to a transparent object, we created images of the glass Glaven where we only colored in those pixels that were 15 CIEDE2000 units or less from the grand mean of the matches that observers made. When paired with the CIEDE2000 color difference metric [76], CIELAB comes closer to being a perceptually uniform space, although still not fully uniform [24], but CIEDE2000 is still the recommended metric for computing JNDs in the CIELAB space, so it is what we use here. The maps were computed with a Go program (v1.13.3) [6] that transformed each test image into the CIELAB space and used a mask to test only those pixels that were filtered by the Glaven. We only computed CIEDE2000 maps for the data coming from the uniform patch experiment with the “proximal match” instructions.

For each test scene that was processed, a new companion image was made that was initially all black. For any pixel in the masked region that passed the test, its original RGB value was placed in the corresponding pixel of the companion black image. All other pixels in the masked region were converted to grayscale using the following formula:

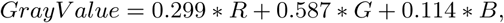

which is the formula used by Go’s color package and which follows the RGB to YCbCr colorspace conversion given in the JFIF specification. This grayscale value was then divided by 1.5, so that the bright intensity of specular highlights would not make them look accidentally colored. The slightly darker gray values also help emphasize the actually colored portions of the map. The resulting grayscale values were then placed in the corresponding pixels in the companion black image. The CIEDE2000 maps are similar to analysis done in Giesel and Gegenfurtner and they give a rough estimate of what might drive the observers’ percepts.

## 3 Results

### 3.1 Matches made for the multi-colored Voronoi background

The average matches of observers for the “proximal match” instructions with a uniform patch, the “dye match” instructions with a uniform patch, and the mean of the flat filter match for the experiments with the multi-colored Voronoi background are shown in Fig. 6. We also show the color of the brightest point forpf the flat filter in Fig. 6. Regardless of the instructions or the matching element, observers’ matches are essentially the same: the chromaticities of their matches are essentially equivalent to the mean chromaticity of the glass Glaven, but the luminance of their matches is consistently higher than the mean. It should be noted that there are indeed differences between the flat filter and uniform patch settings, although they are generally not extreme. This is considered in more detail in the “Discussion” section.

**Figure 6:**
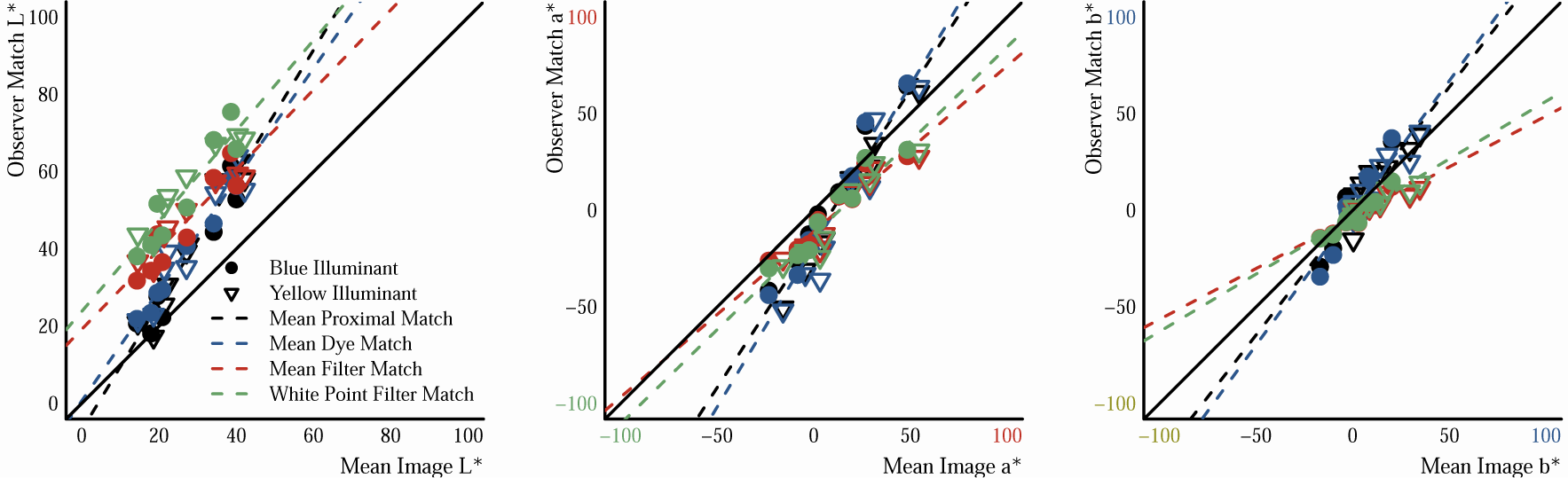
Results of matching experiments with the multi-colored Voronoi background. Each panel shows the location of the resulting match on the three axes of the CIELAB color space: left panel = L*, middle panel = a*, and right panel = b*. The endpoints of the axes in the middle and right panels are colored to indicate the approximate perceived colors along the respective color dimension. The x-axis in each panel represents the mean of the filtered region of the test image (i.e., the region occuppied by the Glaven) for the respective color dimension. The y-axis represents the average match of observers for the same color dimension. Filled dots are matches made under the blue illuminant and empty triangles are matches made under the white illuminant. The color of the symbols denotes the experimental condition: black = proximal match (uniform patch), blue = dye match (uniform patch), red and green = filter match. The difference between the red and green points is whether we plot the mean of the filter match on the y-axis or the color of the brightest point (i.e., the “White Point”) of the filter match. In both cases, the mean and the “White Point” of the filter match are plotted against the mean of the image. The regression lines indicate the general trend of observer settings and can be roughly compared to the unity line to see whether or not observers’ settings track the mean color of the region filtered by the Glaven. In this and all following plots, these regression lines were always fit to the distribution of observer means, not to the grand mean data. Please see “Analysis of observer data” for more info.

In Fig. 6, we have also plotted the color of the brightest point of the flat filter matching element against the mean of the glass Glaven. The are two reasons for this. First, since the filter matching element has spatial variation, there are a number of features that observers could attempt to match to the glass Glaven and the mean does not necessarily have to be one of these features. Second, it is known from work with curved Lambertian objects [34, 81, 80] and glossy objects [36], that when observers are given a uniform patch and asked to set it to the color of the object, they often match it to the brightest region on the object, excluding highlights in the case of the glossy object. This implies that observers might be following a similar strategy here and that regardless of whether they use a uniform patch or a flat filter as the matching element, they will match “brightest” region to “brightest” region. If this were the case, then the green points should considerably differ from the red points in each panel. However, the green points essentially overlap the red points in each panel, so it is unlikely that the “brightest” region of a transparent object is what drives the associated color percept.

If we look at the CIEDE2000 maps in Fig. 7 for the “proximal match” uniform patch instructions, we find that observer matches are close to colors on many different regions of the Glaven stimuli. There is no single specific region that stands out from others. It seems instead that observers are doing their best to squeeze all the variations in the Glaven stimuti into the single color that the uniform patch allows them to set. However, this analysis is only a rough estimation and is not conclusive on its own.

**Figure 7:**
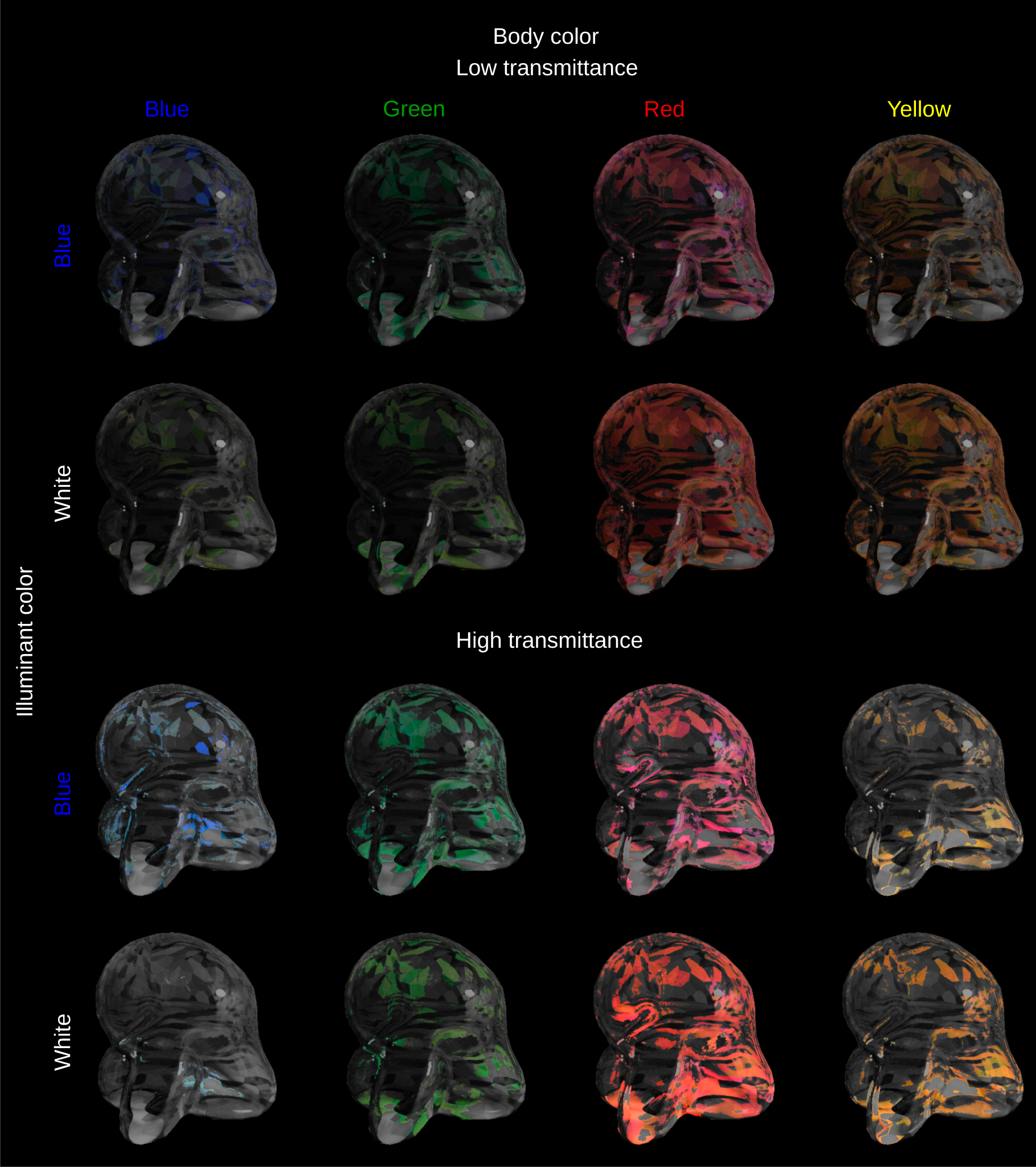
Maps of the glass Glaven stimuli, where pixels are colored in with their original color if that color is located 15 CIEDE2000 units or less from the color of the “proximal match” with the uniform patch matching element (the color in question being the grand mean of observer matches; see “Methods” section for more details). The pattern is essentially the same for the “dye match” instructions. See main text for an interpretation of the results.

### 3.2 Matches made for the white walls scene

In the case of the white walls scene, we only tested the flat filter matching element. The results can be seen in Fig. 8: now, both the chromaticity and luminance of matches correspond closely with the mean chromaticity and luminance of the glass Glaven. The point of this experiment was to test if observers are potentially accounting for the illumination difference between the test scenes and the flat filter renderings. In other words, it tested if observers were performing a kind of color constancy operation. Basically, if we use a scene with a white background and a neutral illuminant, then observers no longer need to discount the illumination difference and the luminance of their matches should come closer to or even directly match the mean luminance of the region filtered by the glass Glaven. We see that this is the case. Taken together, this indicates that observers are performing a color constancy-esque discounting operation that takes into account the fact that the flat filter is rendered on an achromatic background and under a neutral illuminant.

**Figure 8:**
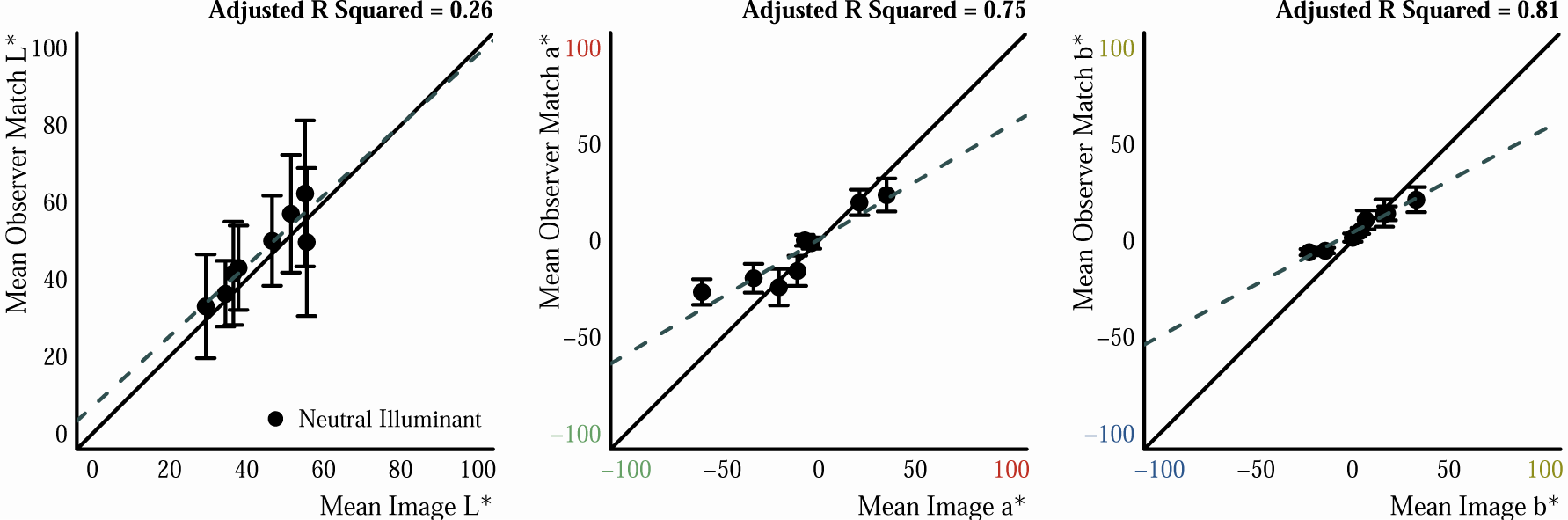
Results of matching experiments with the white background and neutral illuminant. Plotting conventions are the same as those in Fig. 6. See the main text for interpretation of the results.

However, these results are not enough on their own to determine if observers put more weight on discounting the effects of the background or on the effects of the illuminant, since both are confounded in this test. The results for matches with the flat filter and the multicolored Voronoi background were obtained for two different illuminants, so we can gain some perspective from that. In Fig. 9, we plot the results for just that experiment in more detail, where the color of symbols corresponds to the body color of the test Glaven and the lightness of the symbols indicates whether that Glaven had a high or low transmittance. We have zoomed the axes in each panel to ease visual comparison of points. Here, we see that observer matches for the same Glaven (same lightness and color of symbol), but under different illuminants (filled circle or empty triangle), are not always at the same mean color, especially if we focus on the b* dimension (right-most panel). This implies that observers are not always completely discounting the illuminant, but, as we will see in the section below, the mean color is not exactly what observers are matching, so a more rigourous assesment of this statement would require further work.

**Figure 9:**
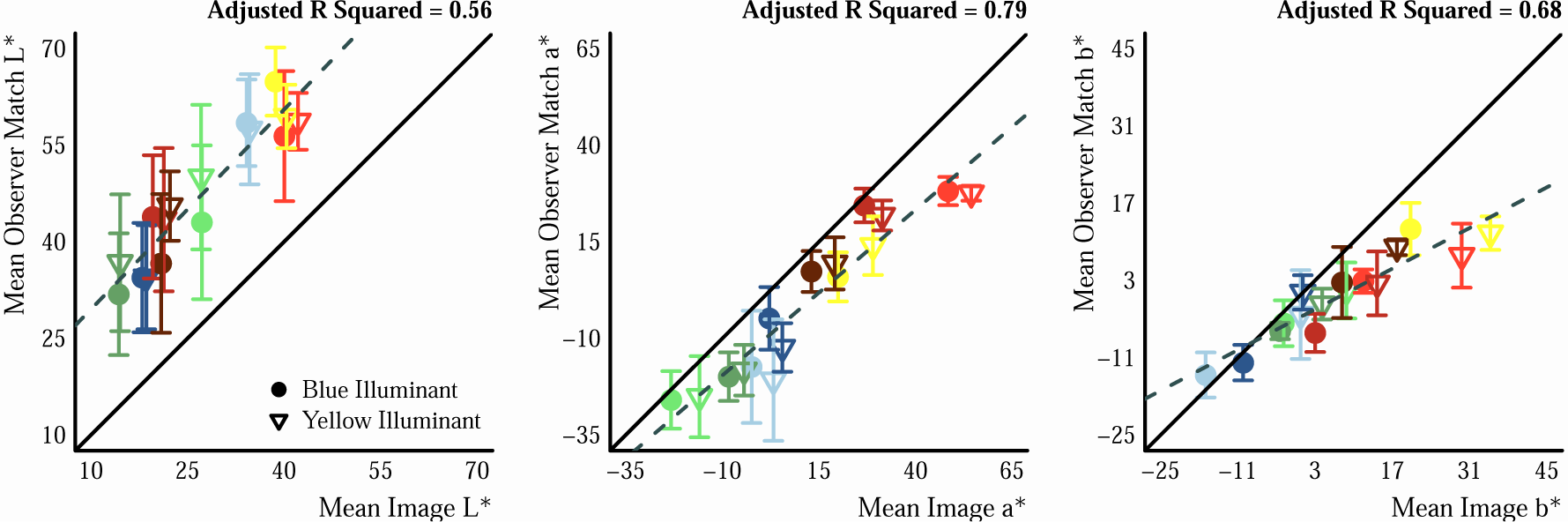
Zoomed results of the experiment with the flat filter matching element and the multicolored Voronoi background. Plotting conventions are the same as those in Fig. 6, except now symbols are colored according to the body color of the test Glaven and the lightness of the symbols corresponds to whether the transmittance of the Glaven was low (darker symbols) or high (lighter symbols). See the main text for interpretation of the results.

### 3.3 Statistics investigated for flat transparent filters

The previous two subsections dealt with the relationship between the mean of the region filtered by the glass Glaven and observers’ matches, where it was found that the mean and the color of the brightest point are not adequate predictors of the matches. However, other important factors that were determined to work for flat filters could certainly translate to curved transparent objects. In the case of the flat filter matching element, there is spatial variation that the visual system could evaluate when detecting transparency and assigning a color to it, and a feature related to this spatial variation could be shared across the flat and curved case. This feature could be what observers are actually matching. Two measures that relate to spatial variation are the ratios of mean cone excitations (RMC) between filtered and unfiltered regions, as suggested by Khang and Zaidi, and the ratios of the standard deviation of cone excitations between filtered and unfiltered regions (RSD), as suggested by Faul and Ekroll, Faul and Ekroll.

In Fig. 10, we compare the RMC and the RSD for the flat filter matching element and those for the test stimulus with the glass Glaven. (See the “Methods” section for details on how these ratios were computed.) We find an initial good correspondence between the RMC and the RSD for the glass Glaven test stimuli and those for the observers’ settings with the flat filter matching element. However, it could be that specific combinations of background colors and filters lead to a breakdown in the correspondence between RMC and RSD. In the case that we chose stimuli that ignore these specific combinations, we ran a series of simulations, where we instructed Mitsuba to render our scene, but for many different combinations of transparency, illuminant, and background distribution. We did this for the lighter and darker transparent distributions, to remain consistent with the experiments detailed above. In particular, we took the four base transparency distributions that were used to render our test scenes with the Glaven (i.e., the spectra that came from the Computational Spectral Imaging group at the University of Eastern Finland [39]) and used those as four different illuminant spectral distributions. We then took these same four base transparency distributions and computed a number of linear combinations of them and used those as the spectral reflectance and spectral transmission distributions for the glass Glaven. To get these new transparency distributions, we took combinations of the base blue, yellow, red, and green distributions and computed linear combinations of them using an equation of the following form:

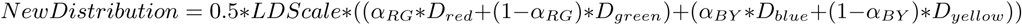

where *LDScale* is a factor that controls whether the resulting distribution gives a lighter or darker body color, *D*_*…*_ is the respective transparency distribution, say the red Munsell transparency distribution, *α*_*…*_ is the multiplication factor that determines the specific linear combination, and the 0.5 scaling factor is used to keep values in the transimission distribution range of [0, 1]. We then independently varied *α*_*RG*_ and *α*_*BY*_ from −1 to 1 in four equally spaced steps and *LDScale* was either 0.68 or 1, like in the experiments described above, which in total would give 32 distributions. We then created four additional background textures, in addition to the one that was already used in the main experiments. Three of them had the same spatial configuration as the multi-colored Voronoi background that we already used (see Fig. 2), but with different color distributions that pushed their means towards red, green, and blue, since the original distribution had a yellow bias. The fourth distribution was an Eidolon-transformed [48] version of a background texture with the blue bias. Briefly, Eidolons apply a locally smooth, but random, spatial deformation to an image (it can be applied separately to different spatial scales), so that the image is distorted but still retains certain features. It can simulate the appearance of objects in the peripheral visual field [48], or when they are under water [21], or what a tarachopic amblyope sees [48]. This Eidolon transformation was done to test if the spatial distribution of the background played a role. The Eidolon-transformed version was also slightly darker. In total, we had five different background textures, since we also used the original background texture with the yellow bias. We then rendered each possible combination of transparent distribution, illuminant distribution, and background texture, giving 640 final images in total.

**Figure 10:**
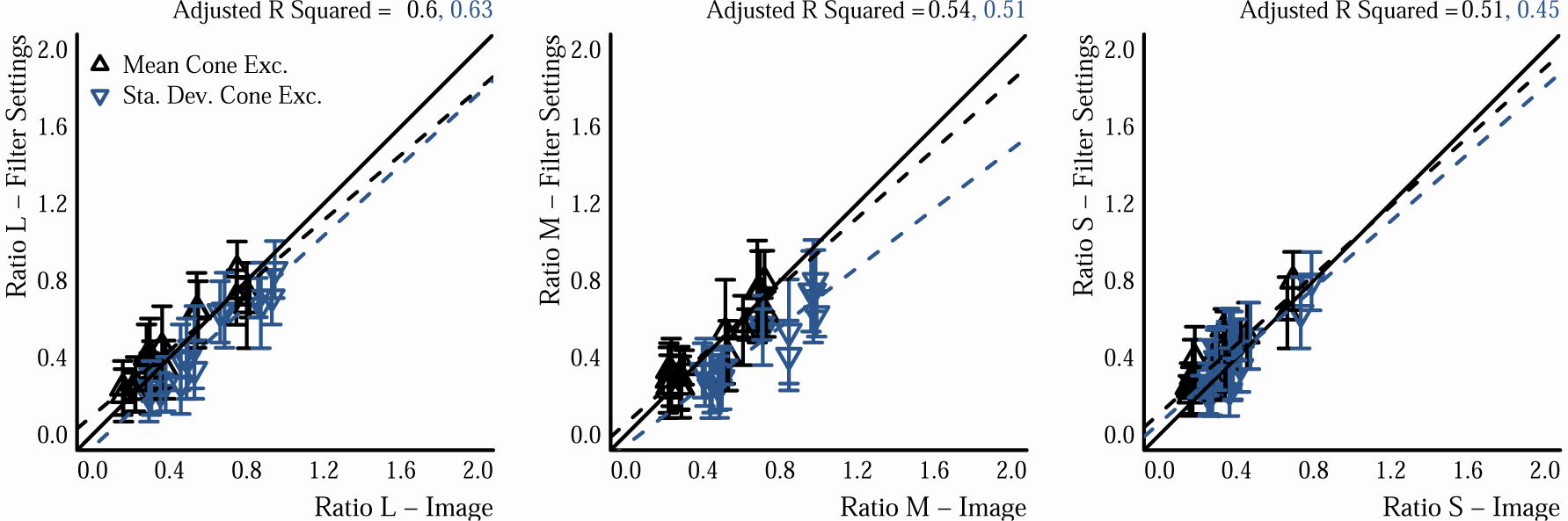
Comparison of RMC and RSD for our test stimuli with the glass Glaven and the matching filter stimulus. For each test scene, we consider each cone type in turn and plot the given statistic for the test scene against the same statistic for the matching filter. Since both statistics are ratios computed from the same units, they are plotted on the same axes as different symbol types (see figure legend). If points lie on the solid unity line, then there is good correspondence between the statistic in the matching filter image and the test scene. The dashed lines show the best fitting linear regression to the data, where their color indicates the associated data points, and the adjusted *R*^2^ for each is shown in the same color above each plot.

After rendering these scenes, we computed the RMC and RSD for them, like described above, and looked at the correspondence between the two statistics. This is shown in panel A of Fig. 11. It can be seen that for some scenes, the correspondence breaks down, especially for the L and M cones. We selected five scenes, shown in panel B of Fig. 11, to see if they could help us determine whether observers equate RMC or RSD when making their filter match. Four of the scenes were chosen such that any pair would have either roughly the same RMC or RSD in the M cone class, while the other statistic would be different. It was too difficult to achieve this criterion across all three cone classes; this is why we arbitrarily applied the criterion to the M cone class. In the end, this was sufficient. The fifth scene was chosen to increase the sample size. Since some of the scenes that break the correspondence are darker overall, we also made brighter versions by rendering the same scene, but with a more reflective background texture: 1.5 times more reflective for the blue, yellow, red, and green background textures, and 9 times more reflective for the Eidolon-transformed background with the blue bias, since it was darker relative to the others.

**Figure 11:**
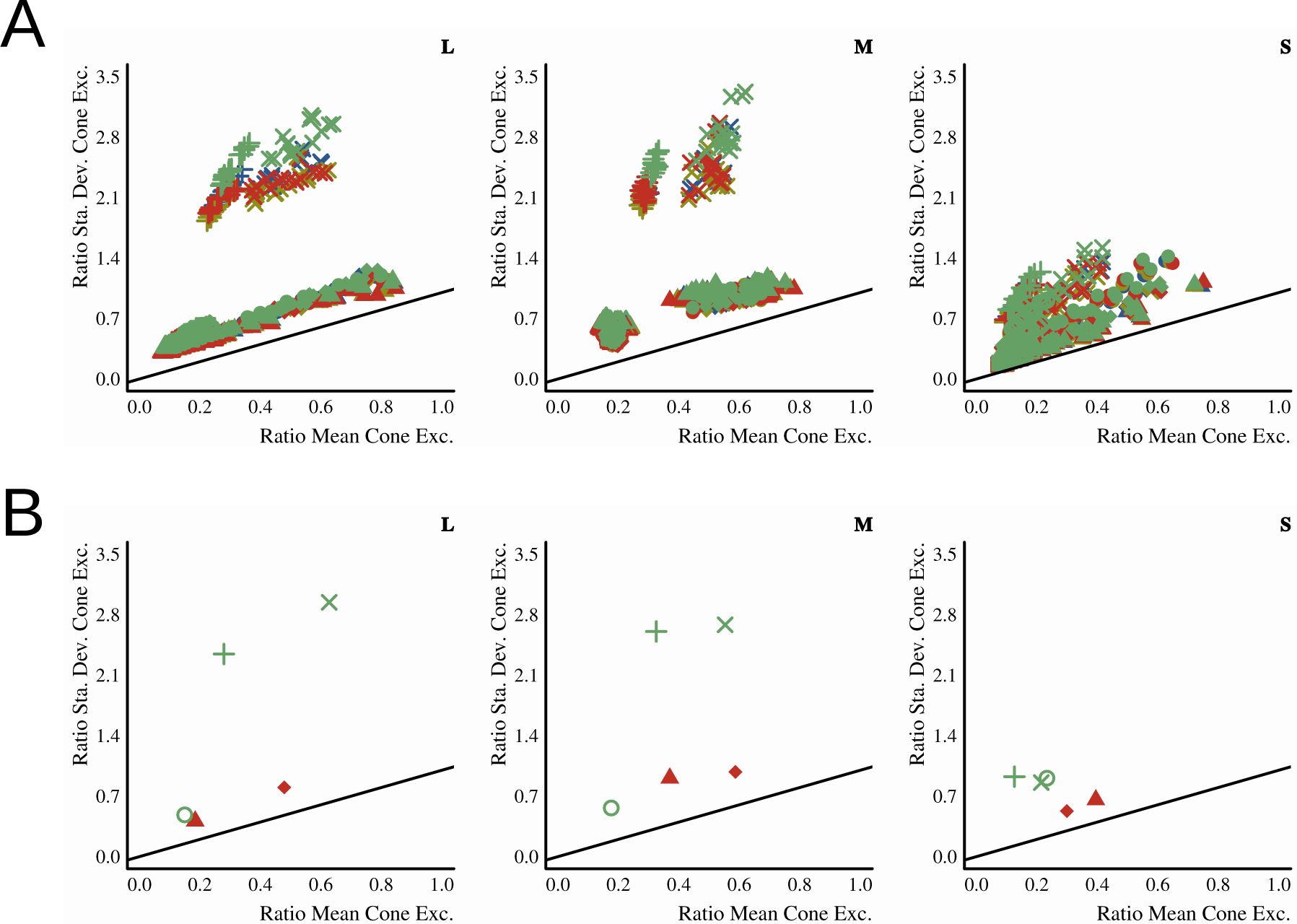
Comparison between RMC and RSD for the simulated scenes described in the main text, computed for each cone class. Panel A) Values for each cone class are in separate subplots. RMC is on the x-axis and RSD is on the y-axis. Each point corresponds to one of the simulated scenes. The colors of the points denote the color of the illuminant, and the plotting symbols simultaneously denote the color bias of the background texture and whether the transparent object had a lighter or darker body color. Lighter body color: open square - blue background, open circle - yellow background, open triangle - red background, open diamond - green background, cross - Eidolon-transformed blue background; Darker body color: filled square - blue background, filled circle - yellow background, filled triangle - red background, filled diamond - green background, x-symbol - Eidolon-transformed blue background. The solid dark line denotes the unity line. Panel B) The values for the five scenes that were chosen for testing.

The ten scenes and the corresponding RMC and RSD values are shown in Fig. 12. In the end, we only presented nine of the ten scenes to the observers and did the data analysis for only those nine scenes, since one of the scenes was too difficult for observers. The scene that was too difficult is shown with a red overlay in Fig. 12. Note that these images are only rough approximations when shown on the internet. The images display better on the Eizo monitor that was used for testing.

**Figure 12:**
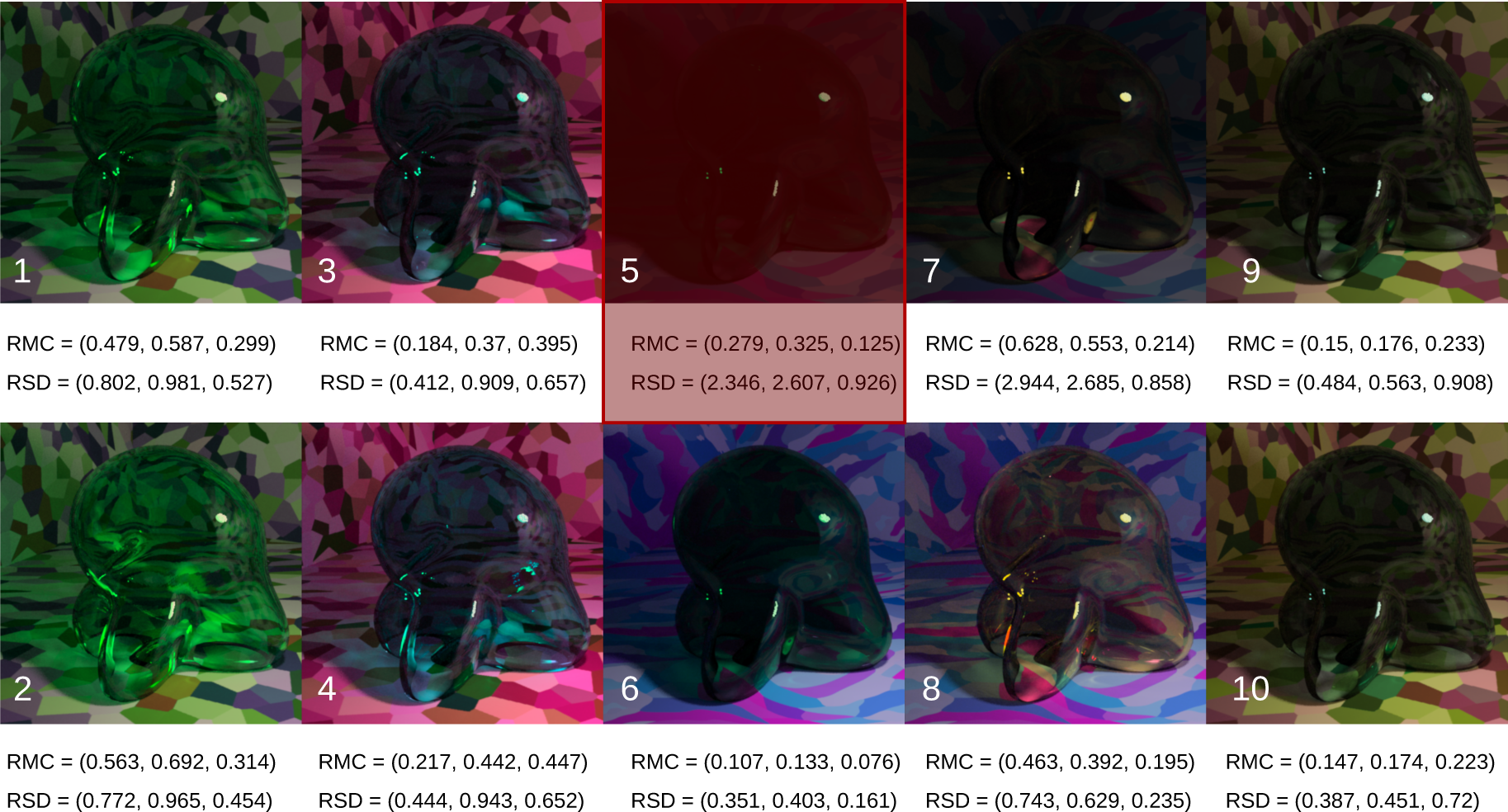
The ten scenes that we used for a more direct comparison of RMC and RSD via our filter matching task. The original scenes that were selected from the simulations are in the top row and the five counterparts with the more reflective background textures are in the bottom row. The images in the third and fourth columns have the Eidolon-transformed blue background texture. RMC and RSD for each scene are shown underneath each image in the following format: (ratio for L cones, ratio for M cones, ratio for S cones). The scene with the red overlay was not tested in the final experiment, since it posed too much difficulty for observers in preliminary testing. Note that these images are only rough approximations when shown on the internet. The images display better on the Eizo monitor that was used for testing. If you are viewing the document on a tablet or laptop, it may also help to tip the display forward or backward a bit.

We then showed these nine scenes to three observers (one of which was the author, RE) and had them make a filter match in the exact same manner as before, except that observers only did 4 repeats for each image, rather than 5, to save a bit of extra time, since observers were rather internally consistent in the previous experiments. Also, this time, after each trial, observers gave a quality rating of their match on an ordinal scale of 1 to 5. It was explained to observers that they should enter 5 if they felt that their match was perfect and that they should enter 1 if they felt that they could not find any satisfactory match, and that they should consider 2, 3, and 4 as equally spaced steps between these possibilities.

The only other major difference from the earlier experiments was that new filter transmission distributions were chosen as the endpoints of the axes that defined the linear combinations observers could make in the flat filter when moving the mouse. We did this because some of these new scenes had glass objects with a body color that was outside the gamut of matches that could be achieved with the selection of transmission distributions used in the experiments described earlier in the paper. The Munsell coordinates that corresponded to the new filters (see “Methods” section for more details) for this specific experiment are shown in Table 4.

**Table 4:** A new set of more saturated Munsell chips whose spectral reflectance distributions that were used in the observer-controlled linear combination that determined the spectral transmittance distribution of the flat filter matching element.

|  |  |
| --- | --- |
| Red | 5RP 4/12 |
| Green | 2.5G 6/12 |
| Blue | 5PB 4/12 |
| Yellow | 2.5Y 7/12 |

Observers’ average quality ratings are shown in Table 5. In panel A of Fig. 13, we show the results for the nine scenes that we tested with the three observers. At first glance, it can be seen that RMC is best at predicting the settings in the S cone channel, and that it is better overall than the RSD, but not perfect.

**Figure 13:**
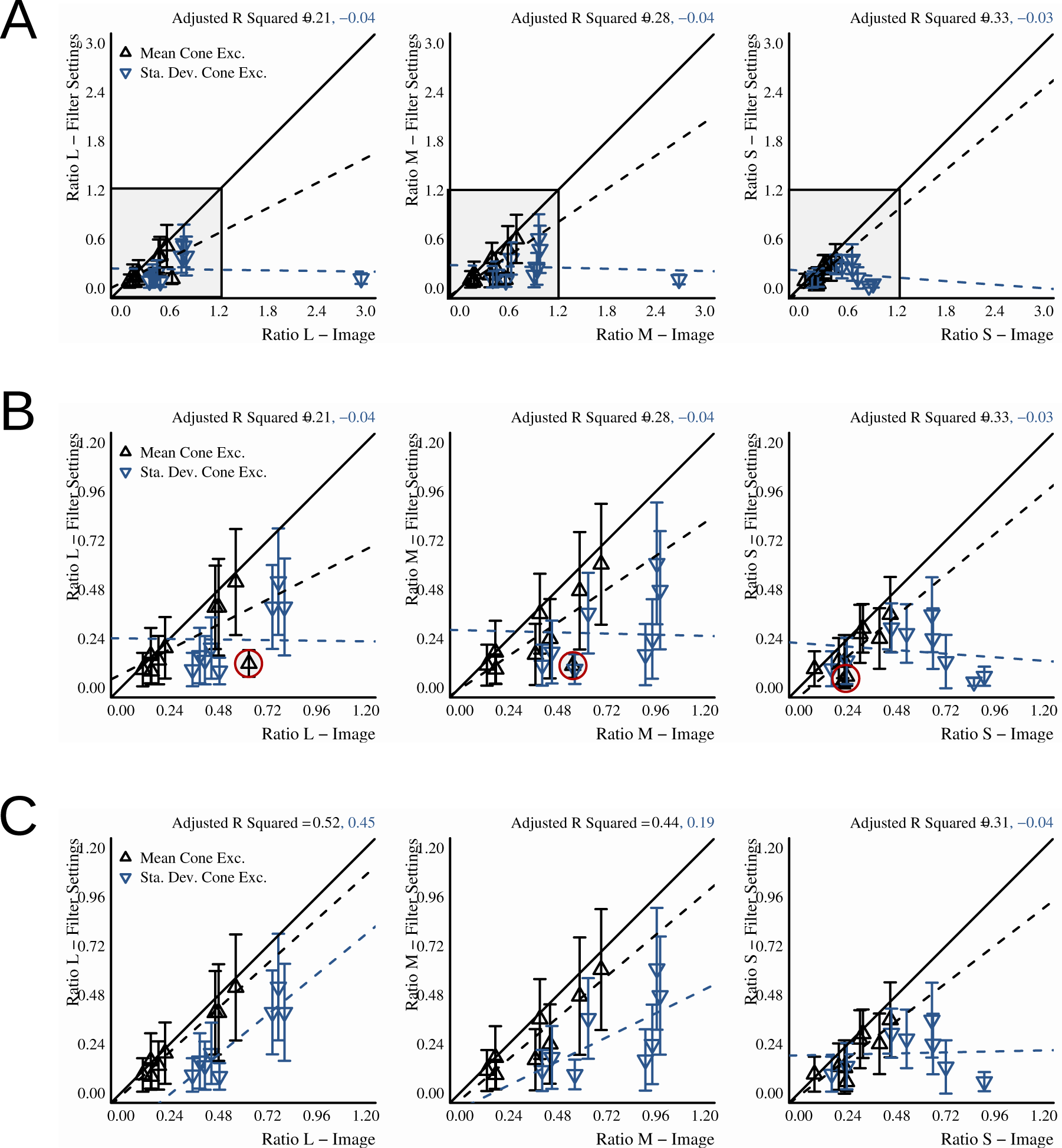
Comparison between RMC and RSD for the nine scenes that are intended to break the correlation between these two statistics (see main text for details). Plotting conventions are the same as in Fig. 11. Panel A) Results for the filter matches of three observers. Panel B) The same data, but zoomed to focus on the shaded, square region in panel A. The best-fit lines are the same as those from panel A, in that they are still fit for the image with very high RSD in panel A. The RMC data for one image is circled in red. This is the same image with the high RSD in panel A. It is an example of an image with a filter setting that has lower RMC than expected given the value of the RMC in the actual test image and the slopes of the best fit lines. Panel C) The same zoom as in panel B, but with the data for the red point removed, since this also corresponded to the image with the lowest quality rating.

**Table 5:** Average quality ratings (±SEM) of observers for their matches to the stimuli that decorrelated the RMC and RSD. The image numbers correspond to the labels in Fig. 12. See the main text for more details.

| Image | Avg. Quality | SEM |
| --- | --- | --- |
| 1 | 4.083 | 0.52 |
| 2 | 3.667 | 0.722 |
| 3 | 4.583 | 0.144 |
| 4 | 4.25 | 0.661 |
| 6 | 4.25 | 0 |
| 7 | 2.333 | 1.041 |
| 8 | 4.25 | 0.661 |
| 9 | 4.25 | 0.5 |
| 10 | 4.667 | 0.289 |

In panel B of Fig. 13, we zoom into the shaded, square region marked in each subplot of panel A. The fitted lines are still for the data shown in panel A, including the image with very high RSD in the L and M cone channels. We see that RMC is doing better overall, but one image, circled in red, is always matched with a filter that has a low RMC setting with respect to the RMC found in the image. This is the same image that has the high RSD value shown in panel A (by design; see explanation of paradigm above), but it is also image 7, which received a low quality rating from observers (see Table 5). Observer filter matches for that image have a very low RSD in comparison to the RSD that is actually in the image. If it is possible that this one image drives the poor trend for RSD, then we can temporarily remove this image from the analysis and look at the results for the eight remaining scenes. That is shown in panel C of Fig. 13. We see that this improves the trend for RMC and RSD, to the point that RMC could be considered a good predictor of the data, whereas RSD is still doing poorly for the M cone channel and especially the S cone channel. However, it is also interesting to actually look at the tested image, its counterpart with the altered background texture, and the average filter settings that observers left in their matches. This is shown in Fig. 14.

**Figure 14:**
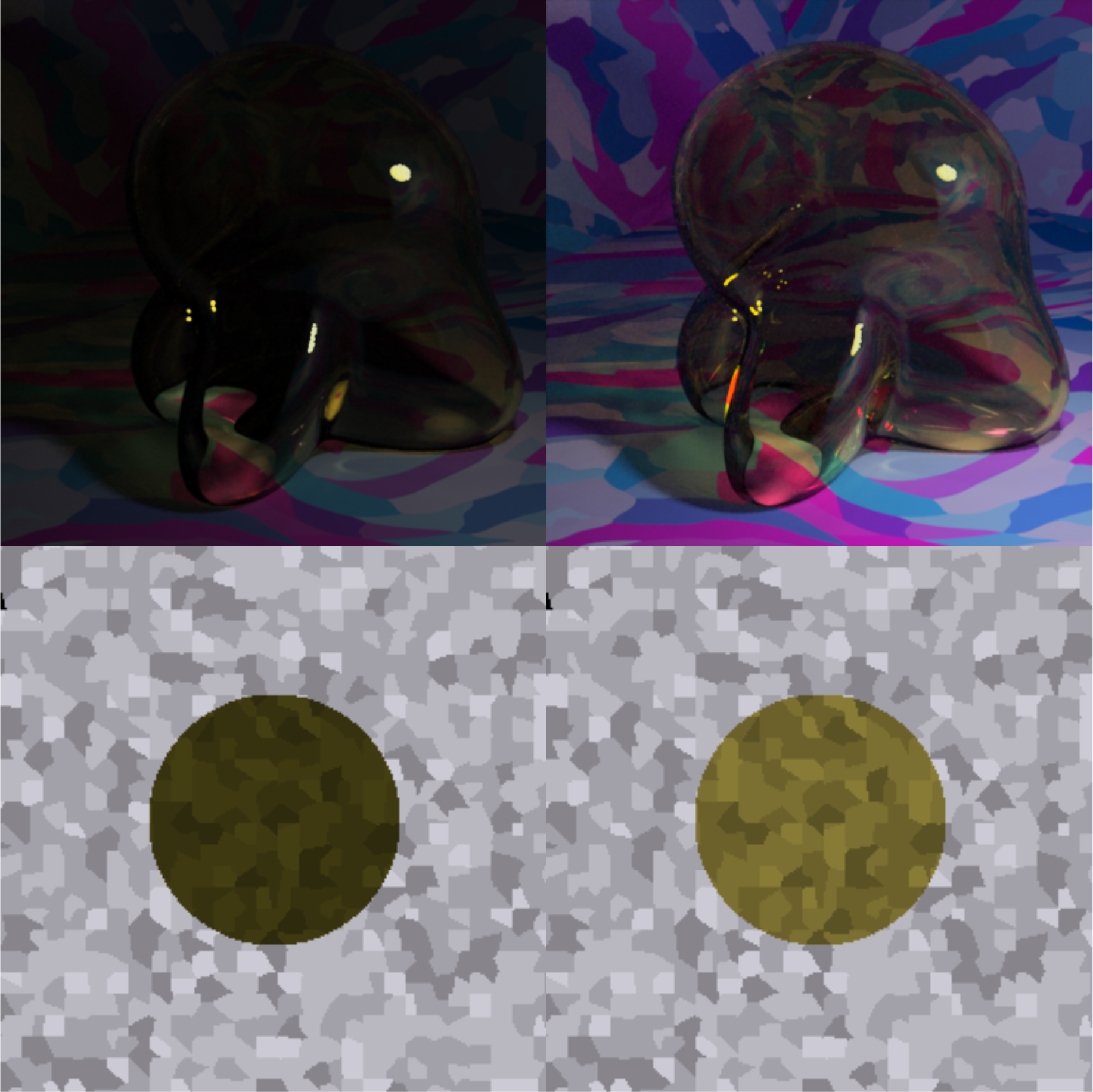
Depiction of the test image whose data are circled in red in panel B of Fig. 13. The image whose data are marked in red is on the top left and its counterpart with the more reflective background texture is on the upper right. The average settings of observers left in the filters when they were satisfied with their match are shown in the bottom row, where the filter on the bottom left is the average match to the image on the upper left and the filter on the bottom right is the average match to the image on the upper right. Note that these images are only rough approximations when shown on the internet. The images display better on the Eizo monitor that was used for testing. If you are viewing the document on a tablet or laptop, it may also help to tip the display forward or backward a bit.

**Figure 15:**
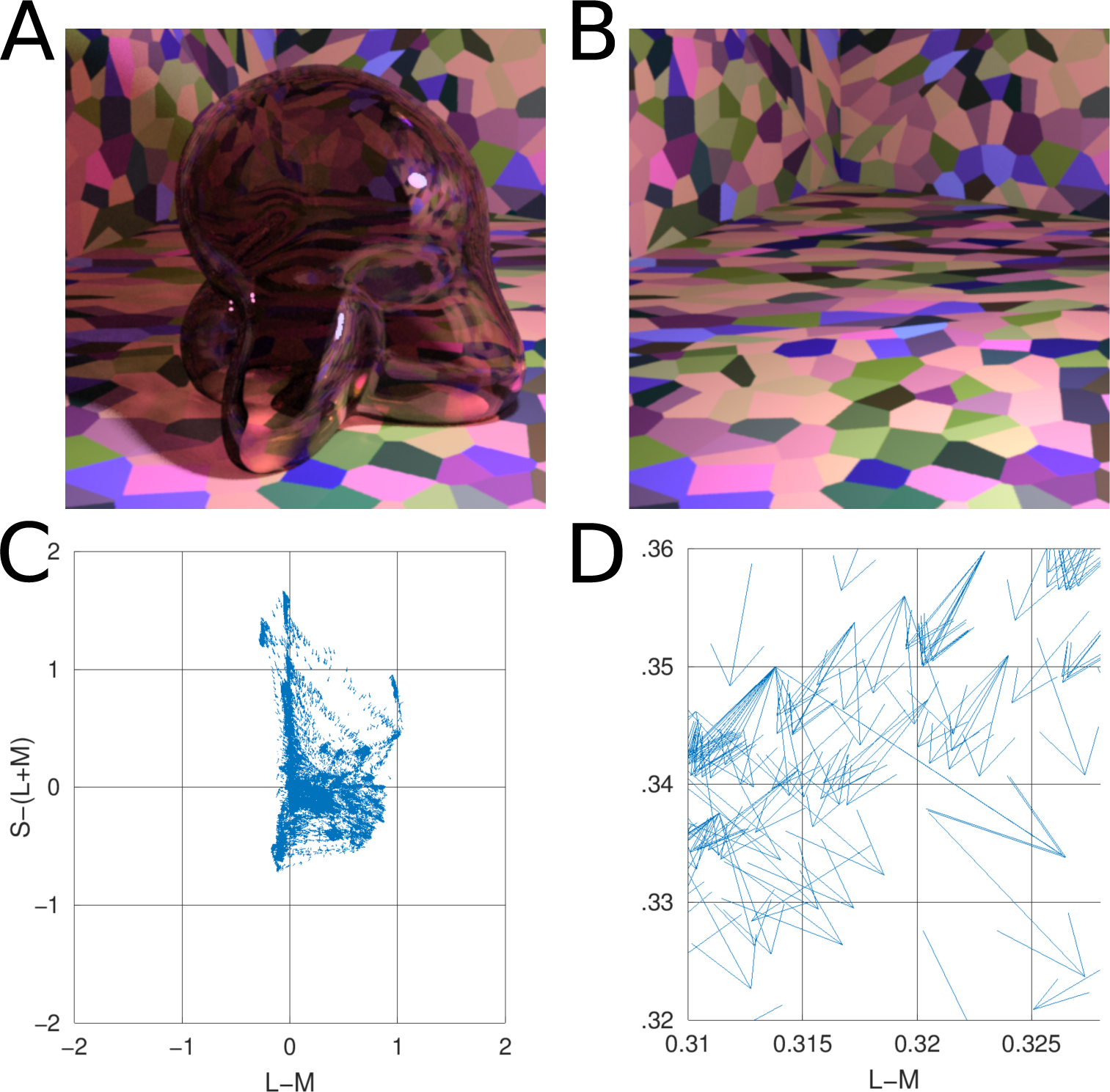
Details of our application of the convergence model to our stimuli. Panel A) The same scene with the glass object as shown in Fig. 1 of the “Introduction” section in the main text. Panel B) This same scene with the glass object removed. Panel C) The vector field that connects the color of pixels in panel B to their filtered counterparts in panel A. Only the pixels that are actually filtered by the glass Glaven are included in the vector field. The bases of vectors are the colors from panel B and the tips are their filtered counterparts from panel A. Panel D) A close-up zoom of a region of the vector field shown in panel C. One can see that many vectors are assigned to the same point in color space. Please note that for presentation purposes the scaling of the vectors is arbitrary, since we plotted the data with MATLAB’s quiver() function.

In Fig. 14, we see that although observers gave a low quality rating for the filter matches to the glass object in the scene on the upper left, they actually make a setting that is reasonable: what appears to be a dark yellow-green glass is matched to a dark yellow-green filter. We show the counterpart image with the more reflective background texture on the right to show that observers make a similar match in terms of hue when the intensity of the scene is greater overall.

This indicates observers might just be hesistant when making a match under darker conditions and although they are making a reasonable match, they just do not trust themselves and put a low quality rating. With a bit of confidence training, observers might put higher quality ratings for such images and then future investigations of these types of images, where both RMC and RSD do not predict the matches that observers make, might provide further insight into the mechanisms that determine the color of a transparent object.

From these simulations and these data, we find that observers’ settings do diverge when RMC and RSD are de-correlated in test images, but in a complicated way. The data support RMC over RSD as a likely image statistic that observers use to assign color to a transparent object, yet there are images that contend with this conclusion, such as the red point in panel B of Fig. 14. These images could indicate that there is another statistic which trades-off with RMC for determining the color of a transparent object. Future work will need to be done with images of this type.

## 4 Discussion

Our goal was to understand what determines the perceived color of a transparent 3-D curved object, and we have found further evidence that the percept is driven by the ratios of the mean cone excitations between the filtered and unfiltered regions of the image (RMC), as suggested by Khang and Zaidi. However, one of our results suggest that additional statistics might be at play or might trade-off with the RMC, especially since there are more factors at play for curved, transparent objects.

Interestingly, regardless of whether they use a uniform patch or a flat filter as the matching element, the mean colors of observer matches are almost the same, but still different. When using the uniform patch, observers are making a very asymmetric match between objects in two different modes of perception [41, 10]. The patch appears like a flat Lambertian surface or a piece of paper, while the glass Glaven is curved and has a body color and various physical properties that the patch cannot emulate. This could explain the discrepancy seen in Fig. 6. Although, we should point out that observers never stated any difficult with any of the instructions nor with either matching element and all observers finished the experiments in roughly 45 minutes.

If we turn our attention to the flat filter matching element, then we find that the mean color of the matched filter is almost the same as the mean color of the glass Glaven itself, while the luminance setting is still offset, even more so than with the uniform patch. This is interesting because there is reduced discrepancy between modes of perception for the curved, glass Glaven and a flat transparent filter. In other words, to match a piece of glass to a curved glass object is probably easier than to match a piece of paper to the same curved glass object. The point is that it does not necessarily need to be the mean color that observers match and the reduced discrepancy in modes of appearance can make it easier for observers to match the statistic that is actually relevant.

Essentially, we find that the RMC is the best predictor to date of the matches that observers make. It is important to point out that the RMC inherently implements a discounting operation, by comparing “background” colors to “foreground” colors (i.e., colors in a region of interest), which helps it be the best predictor across different background and illuminant scenarios. The relationship between color constancy and ratios of cone absorptions, computed in a fashion similar to the RMC, is covered elsewhere [87], but since transparent filters share some characteristics with spotlight illuminants [45, 18, 47], it is sensible that the visual system would reuse strategies. However, as stated in the “Matches made for the white walls scene” section, whether observers place more focus on discounting the effects of the background colors or the effects of the illuminant is still an open question for curved, transparent objects. Given this, it may not be too much of a stretch to say that transparent objects are like “embodied light”. Although, it is necessary to remember that transparent objects still have a number of their own individual quirks that distinguish them and there may be more than the RMC at play for 3-D, curved, transparent objects.

The use of a flat filter as a means to determine the perceived color of a transparent object does raise some interesting questions. First, it is important to keep in mind that it is not the case that the matches with the uniform patch could be “wrong” and that those with the flat filter could be “more accurate” or “right”. They are just different and observers are using different strategies to account for those differences. The original intention of the uniform patch matching element was to see if observers can reduce the appearance of a transparent object to a single point in color space (i.e., a single color), since we certainly see a red glass object as “red” with a specific and distinct color. Regardless of whether observers use a uniform patch or a flat filter, they make matches that to us (the authors) look sensible, but the actual settings do differ in their color properties. This is reminiscent of the comment by Xiao and Brainard that there may not actually be a single well-defined color for all three-dimensional objects.

On a related note, Richards et al. have found that if one restricts themselves to the Metelli model of flat transparent filters, then the model can state that certain filter and background combinations can lead to a filter with a color that is not physically realizable in the standard CIE1931 xyY colorspace. Considering this, it may be the case that our stimuli do not push the limits of transparent colors, where observers might not be able to make a match with a uniform patch. On the other hand, the CIE1931 xyY colorspace may not be the appropriate space for representing “transparent colors”. All of this starts to bring in a chicken-and-egg question. Consider that the flat filter is essentially a “uniform patch” for transparent objects: it has none of the specular highlights, shadows, or caustics that appear with the glass Glaven, much like the uniform patch has no shadows and none of the curvature that a typical Lambertian object would have. Yet, to actually perceive the filter as transparent, it needs to be placed over a variegated background, like the achromatic Voronoi background that we used, so that the color of the filter is tied in some way to the pattern which it covers. If we instead place the filter over a multi-colored background, then we quickly notice that we essentially return to our original problem: what is the specific color of a transparent flat filter? While matching a uniform patch to a flat filter is certainly easier than matching a uniform patch to a curved, glass object and it may actually be quite informative about the color of the flat filter, it would be too much of a stretch to match a filter to the glass Glaven, for example, and then match a uniform patch to the filter and conclude that with the final setting of the uniform patch, we have found the “specific color” that corresponds to the glass Glaven (or even that of the flat filter!). The point of this is to emphasize that colors of different materials do exist in their own individual classes, much as Beck and Katz suggested, but it seems that observers can make mappings between the different classes, regardless.

Our results stand in contrast with earlier work on the color of Lambertian and glossy objects, such as that by Giesel and Gegenfurtner, Toscani et al., Toscani et al., and Granzier et al.. This body of work found that when provided with a uniform patch matching element, observers match to the most luminant region on the body of a Lambertian object and to the most luminant region (excluding the highlights) on the body of a glossy object. Xiao and Brainard came to similar conclusions when they found that observers compensate for the specular highlights on glossy three-dimensional objects, when they are asked to change the color of a matte sphere to match that of a glossy test sphere. However, in these cases, the results are more readily interpretable. The modes of appearance for a uniform patch and a Lambertian object are similar in the same way that the flat filter is similar to the glass Glaven, so the conclusion that the color of a Lambertian object is determined by its most luminant region makes sense and is a good strategy, since the visual system will have a strong signal to work with from the most luminant region. In the case of glossy objects, that observers are matching to the most luminant region, not including the highlights, indicates that they realize that the highlight color is not directly informative about the color of the glossy object and that they can compensate for the effects of specular reflections.

We suspect that the different results that we find with the flat filter could be due to transparent objects being an example of volume colors [41, 63, 10].

## 5 Conclusion

Observers readily answer the question “what is the color of this transparent object?”, regardless of whether the matching element is a uniform patch or a flat transparent filter. In both cases, the chromaticities of their matches are close to the mean chromaticity of the glass Glaven, but the luminance of their matches is consistently offset from the mean luminance. This offset appears to be the result of a color constancy-esque discounting operation. For the flat filters, we can say that the ratios of the mean cone excitations (RMC) between the filtered and unfiltered regions is more exactly what is matched. However, some of our results also indicate that other sources of information could be at play, which might correlate with or trade-off with the RMC and further work will be needed to determine that.

## 6 Acknowledgments

We would like to thank the following for productive discussions: Qasim Zaidi, Karl Gegenfurtner, Arthur Shapiro, Matteo Toscani, Matteo Valsecchi, and Roland Fleming. Thanks to Anya Hurlbert for suggesting that we include the “White Point” statistic and Christoph Witzel for suggesting that we perform the white walls experiments. Funding was provided by the Alexander von Humboldt Foundation in the framework of the Sof’ja Kovalevskaja Award endowed by the German Federal Ministry of Education and Research.

## 7 Supplemental Material

### 7.1 Convergence model analysis details

Here, we detail the application of the convergence model [19, 14, 20] to the stimuli in our paper. Consider Supplemental Fig. 15. We show again, in panel A, one of our stimuli with a glass Glaven in a multicolored room. Panel B also shows how this scene looks when the glass object is absent. Panel C depicts a vector field which shows how the colors from the region filtered by the glass Glaven are transformed when going from the image in panel B to the image in panel A. The bases of the vectors correspond to the colors from the image in panel B and the tips correspond to the colors from the same pixels after they have been filtered by the Glaven in panel A. Panel D shows a close-up zoom of a region in the center of Panel C. It details how the vector field that characterizes the color change between the filtered and non-filtered scenes (i.e., the images in panel A and B) can have many vectors assigned to the same point in the chromaticity plane. For the initial Affine mapping analysis discussed in the main text, we performed the following procedure in MATLAB (R2018b; Mathworks, Inc.; Natick, Mass., USA):

1. Use the same mask for both images to extract the pixels that are directly filtered by the transparent object.
2. Convert the RGB colors of these pixels to the three-dimensional MB-DKL and CIELAB color spaces (see the “Methods” section in the main text of the paper for descriptions of these spaces).
3. Find an Affine transformation that best maps the unfiltered colors from the masked region of the image in panel B to their filtered counterparts from the image in panel A.
4. Repeat the Affine transformation search, but ignoring luminance (i.e., all colors are projected into the mid-gray isoluminant plane and only chroma is considered). This emulates the original investigations of the convergence model.
5. Calculate the RMSE between the actual filtered colors in the masked region in panel A and the results of applying the best-fitting Affine transformation to the unfiltered colors. RMSE should be relatively close to zero relative to the scaling of the axes in the space of consideration.

We repeated this same analysis three times, using three masks: a mask that captures the whole body of the transparent object and that includes the specular highlights and specular reflections; a mask that excludes the specular highlight; and another mask that excludes specular reflections and specular highlights (see the “Methods” section in the main text of the paper for details on how the specular reflection regions were computed). A table with the average RMSE results from this analysis is shown in Table 1 of the “Introduction” section in the main text.

Since the original convergence papers spoke specifically of “convergence”, we decided to also apply the vector calculus operation of divergence [75] to the colors filtered by the glass Glaven as an alternative formulation of the model. To do this, we computed the vector difference in color space between the filtered and unfiltered colors at each pixel, giving a vector field that showed how each pixel in the masked region of panel B of Supplemental Fig. 15 was transformed to the color in panel A. The vector field in MB-DKL space for the example in panel A is shown in panel C: the bases of the vectors are the unfiltered colors from panel B and the tips of the vectors are their filtered counterparts from panel A. Since the walls of the room in our stimulus have broad regions that are all the same color, it was found that many pixels from panel B map to the same point in color space. However, because the curved transparent object has a different optical thickness for filtering at each point in the image and because specular reflections, caustics, and shadows are also in the filtered region, many of these pixels that initially map to the same point eventually spread out in color space after filtering and map to many different points. Panel D of Supplemental Fig. 15 shows a zoomed region of panel C to highlight this. However, this pattern is not conducive to divergence calculations, since it is a one-to-many mapping, resulting in a vector field that is not continuously differentiable in the format that is expected by the classical divergence formula.

To account for this, we created a basic interpolation approach that went through the bounding box around the cloud of vectors, at equally spaced intervals along the axes. At each point, any vectors within a small box centered at the point of interest, whose side length was equal to the step size of the interpolation algorithm, were averaged and that average vector was saved into a separate array at that same point. If no vectors were within the box, then a vector of length zero was saved at the point in the separate array. The result was a vector field that was very similar to the original, but with each point having only one vector associated with it. The result of this interpolation procedure for the scene considered in Supplemental Fig. 15 is show in panel C of Fig. 1 in the “Introduction” section of the main text of the paper. The divergence() function of MATLAB was then applied to this field. Please note that this interpolation method and the resulting divergence analysis were only computed for the chromaticity components of the vector fields, excluding luminance (i.e., after the vectors were projected into the isoluminant plane).

